# Microbial Nitrogen Metabolism in Chloraminated Drinking Water Reservoirs

**DOI:** 10.1101/655316

**Authors:** Sarah C Potgieter, Zihan Dai, Stefanus N Venter, Makhosazana Sigudu, Ameet J Pinto

**Author notes:** corresponding author: Dr Ameet J Pinto, *Email address.

## Abstract

Nitrification is a common concern in chloraminated drinking water distribution systems. The addition of ammonia promotes the growth of nitrifying organisms, causing the depletion of chloramine residuals and resulting in operational problems for many drinking water utilities. Therefore, a comprehensive understanding of the microbially mediated processes behind nitrogen metabolism together with chemical water quality data, may allow water utilities to better address the undesirable effects caused by nitrification. In this study, a metagenomic approach was applied to characterise the microbial nitrogen metabolism within chloraminated drinking water reservoirs. Samples from two geographically separated but connected chloraminated reservoirs within the same drinking water distribution system (DWDS) were collected within a 2-year sampling campaign. Spatial changes in the nitrogen compounds (ammonium (NH_4_^+^), nitrites (NO_2_^−^) and nitrates (NO_3_^−^)) across the DWDS were observed, where nitrate concentrations increased as the distance from the site of chloramination increased. The observed dominance of *Nitrosomonas* and *Nitrospira*-like bacteria, together with the changes in the concentration of nitrogen species, suggests that these bacteria play a significant role in contributing to varying stages of nitrification in both reservoirs. Functionally annotated protein sequences were mined for the genes associated with nitrogen metabolism and the community gene catalogue contained mostly genes involved in nitrification, nitrate and nitrite reduction and nitric oxide reduction. Furthermore, based on the construction of Metagenome Assembled Genomes (MAGs), a highly diverse assemblage of bacteria (i.e., predominately *Alpha*- and *Betaproteobacteria* in this study) was observed among the draft genomes. Specifically, 5 MAGs showed high coverage across all samples including two *Nitrosomonas, Nitrospira, Sphingomonas* and a *Rhizobiales*-like MAGs. The role of these MAGs in nitrogen metabolism revealed that the fate nitrate may be linked to changes in ammonia concentrations, that is, when ammonia concentrations are low, nitrate may be assimilated back to ammonia for growth. Alternatively, nitrate may be reduced to nitric oxide and potentially used in the regulation of biofilm formation. Therefore, this study provides insight into the genetic network behind microbially mediated nitrogen metabolism and together with the water chemistry data improves our understanding nitrification in chloraminated DWDSs.

## 1. Introduction

Disinfection of the drinking water in some DWDSs is often considered key in the management of microbial growth and the maintenance of water quality in most parts of the world, with the exception of a few countries in Europe where microbial growth in drinking water distribution systems (DWDSs) can be managed through nutrient limitation (van der kooij *et al*., 2002; Hammes *et al*., 2008). Chlorine and chloramine have long been successfully used to control microbial growth within DWDSs and although chlorination (primary disinfectant) is successful at initially reducing bacterial growth, distribution system management often now includes chloramination as a secondary disinfectant. Chloramines are typically used to provide disinfectant residuals when free chlorine residuals are difficult to maintain. Chloramines show greater stability as compared to chlorine in the DWDS over long distances, increased efficiency in reducing biofilm growth, and they also produce lower concentrations of regulated disinfection by-products (Norton and LeChevallier, 1997; Vikesland *et al*., 2001; Regan *et al*., 2003; Zhang *et al*., 2009).

However in chloraminated systems, the introduction of ammonia provides an alternative source of nitrogen and growth substrate for ammonia-oxidising microorganisms (AOM), either due to the presence of excess free ammonia or through ammonia released due to chloramine decay (Regan *et al*., 2003; Zhang *et al*., 2009). This promotes the growth of nitrifying bacteria and archaea, leading to increased nitrification (Belser, 1976; Nicol and Schleper, 2006). Nitrification is an essential process in the biogeochemical nitrogen cycle and links the aerobic and anaerobic pathways of the nitrogen cycle by delivering nitrite and nitrate as electron acceptors for dissimilatory nitrate reduction, denitrification, respiratory ammonification, and anaerobic ammonia oxidation (Kraft *et al*., 2014; Koch *et al*., 2015). Traditionally microbial nitrification was considered as a two-step process: firstly, ammonium (NH_4_^+^) is oxidised to nitrite (NO_2_^−^) by chemolithoautotrophic ammonia-oxidizing bacteria and archaea (AOB and AOA, respectively) (van der Wielen *et al*., 2009) and secondly nitrite is oxidised to nitrate (NO_3_^−^) by chemolithoautotrophic nitrite-oxidising bacteria (NOB) (Wolfe *et al*., 1990; Cunliffe, 1991; Francis *et al*., 2005). NOB are often the principle biological source of nitrate, which is not only an important source of nitrogen for other microorganisms but can also serve as an electron acceptor in the absence of oxygen. In contrast to AOB/AOA and NOB, complete ammonia oxidising bacteria (i.e., comammox) can completely oxidise ammonia to nitrate (Daims, *et al*., 2015; Pinto *et al*., 2015; van Kessel *et al*., 2015). Unlike AOB, AOA, and NOB, which are phylogenetically diverse, all known comammox bacteria, belong to the genus *Nitrospira* (Phylum: Nitrospirota) (Daims, *et al*., 2015; Pinto *et al*., 2015; van Kessel *et al*., 2015; Fowler *et al*., 2018; Palomo *et al*., 2018). Furthermore, reciprocal feeding has been described where NOB from the genus *Nitrospira* initiate nitrification by supplying ammonia oxidisers lacking urease and/or cyanase with ammonia from urea or cyanate (Lücker *et* al., 2010; Koch *et al*., 2015; Palatinszky *et al*., 2015).

Bacterial nitrification in the DWDS causes depletion of chloramine residuals and disinfection decay. The resulting formation of nitrite in the system is problematic as it can rapidly decrease free chlorine and is also further oxidised leading to an accelerated decrease in residual chloramine (Wolfe *et al*., 1990; Cunliffe, 1991). In addition, due to its toxicity, the regulated concentration of nitrite is typically very low. While ammonia, nitrites and nitrates can serve as an energy source for AOB and NOB (Kirmeyer *et al*., 1995; Pintar and Slawson, 2003), the loss of chloramine residuals can also lead to heterotrophic bacterial growth and biofilm accumulation, which potentially causes operational problems for many drinking water utilities (Kirmeyer *et al*., 1995; Norton and LeChevallier, 1997; Pintar and Slawson, 2003).

In a previous study by Potgieter *et al*. (2018), *Nitrosomonas* spp. were observed to be dominant in the chloraminated sections of the DWDS, suggesting that ammonia oxidation and potentially nitrification may be important processes in this DWDS. Therefore, the principle goal of this study was to understand the metabolic potential of microbial communities that might impact the fate of nitrogen in a chloraminated DWDS. Here, the organisms and genes involved in the nitrogen cycle, using a genome resolved metagenomics approach, was investigated. The use of shotgun metagenomic sequencing allowed us to: (i) overcome primer bias, (ii) assemble large operons through *de novo* assembly and (iii) to pinpoint functions to genomes through binning, which can then be phylogenetically identified. Therefore, using this approach, the study aims to explore nitrogen metabolism in chloraminated drinking water reservoirs by (i) investigating the taxonomic profile of the microbial community and the specific genes involved in nitrogen metabolism, (ii) identifying the processes that can drive nitrogen transformation in chloraminated drinking water, and (iii) identifying the role of dominant reconstructed draft genomes in nitrogen metabolism.

## 2. Materials and methodology

### 2.1 Site description and sample collection

Sampling was conducted at two geographically separated but connected chloraminated reservoirs within a large South African DWDS previously described by Potgieter *et al*. (2018). Briefly, the process for treating surface water includes coagulation with polymeric coagulants, flocculation, sedimentation, pH adjustment with CO_2_ gas followed by filtration (rapid gravity sand filters) and finally initial disinfection with chlorine. Filter effluent is dosed with chlorine to achieve total residual chlorine concentrations varying between 1 and 1.5 mg/L at the outlet of the drinking water treatment plant (DWTP). Chlorinated drinking water is then dosed with chloramine (0.8 to 1.5 mg/L) at a secondary disinfection boosting station approximately 23 km from the DWTP. Here, monochloramine residuals vary seasonally between 0.8 and 1.5mg/L. Within the chloraminated section of the DWDS, the first of the two reservoirs (RES1) sampled is located approximately 32 km from the secondary disinfectant boosting station. The second reservoir (RES2) is located approximately a further 88 km downstream from the first reservoir (Fig. 1). Samples were collected within 2 years (October 2014 to September 2016). Further details on a range of chemical parameters, including temperature, disinfectant residual concentrations (i.e., free chlorine, total chlorine, and monochloramine) and nitrogen species concentrations (i.e., ammonium, nitrite and nitrate) were obtained from the utility (Table S1A and S1B).

**Fig. 1:**
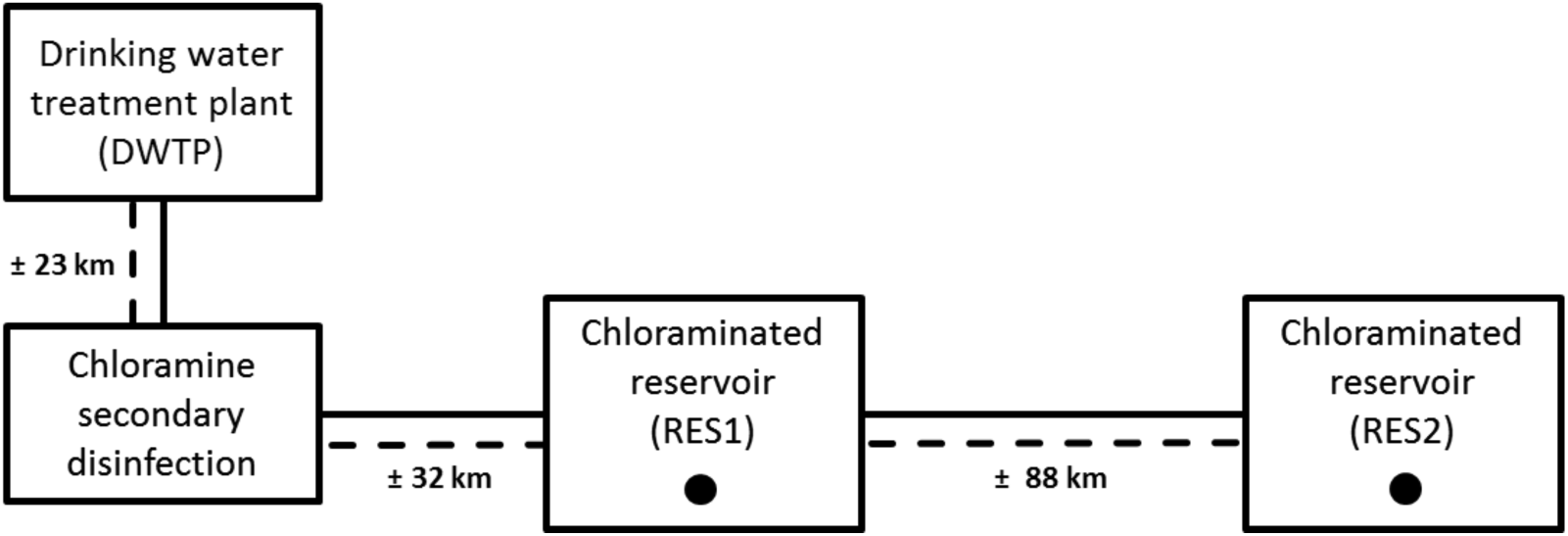
Simplified schematic showing the layout of the DWDS and the sample locations (RES1 and RES2 indicated in the figure as black circles). Approximate distances between locations indicated in the figure as the dotted line.

### 2.2 Sample processing

Bulk water samples were collected in 8L sterile Nalgene polycarbonate bottles and transported to the laboratory on ice where they were kept at 4 °C for 24 to 48 hours until further processing. Samples were filtered to harvest microbial cells by pumping the collected bulk water through STERIVEX™ GP 0.22 μm filter units (Millipore) using a Gilson^®^ minipuls 3 peristaltic pump. The filters were kept in the dark and stored at −20 °C until processing and DNA extraction. A traditional phenol/chloroform extraction method optimised by Pinto *et al*. (2012) modified from Urakawa *et al*. (2010) was used for the isolation of DNA from cells immobilised on filter membranes. Following extraction, 8 samples from RES1 and 10 samples from RES2 were selected for shotgun metagenomic sequencing (Table S2).

### 2.3 Metagenomic sequence processing, *de novo* assembly, functional annotation, and reference mapping

Paired end sequencing libraries were prepared using the Illumina TruSeq Nano DNA Library Preparation kit. Metagenomic sequencing was performed using the Illumina HiSeq 2500 sequence platform at the Agricultural Research Council – Biotechnology Platform (ARC-BTP), Gauteng, South Africa, resulting in 250 nt paired–end reads (13,267,176 ± 3,534,751 reads per sample). Prior to assembly, the metagenomic reads were subject to adaptor removal and quality filtration using Trimmomatic (Bolger *et al*., 2014) with a minimum sliding window quality score of 20 and reads shorter than 100 bp were discarded. Following quality filtering, the level of coverage of each metagenome was assessed using Nonpareil, a statistical program where read redundancy is used to estimate coverage (Rodriguez and Konstantinidis, 2013). Prior to assembly, metagenomic reads were pooled and *de novo* assembly of quality trimmed reads into contiguous sequences (contigs) followed by scaffolding using metaSPAdes assembler version 3.9.0 (Nurk *et al*., 2017) was performed with kmers list of 21, 33, 55, 77, 99, 127. The resulting assembly consisted of 1,007,176 scaffolds (> 500 bp) and an N50 and L50 of 1638 bp and 42230 bp, respectively. Reads were mapped to the scaffolds greater than 500 bp, bam files were filtered to retain mapping reads (samtools view –F 4), and the number of reads mapping to the scaffolds in the metagenomics assembly were counted using awk script. An average of 13,092,168 ± 3,561,160 reads per sample (> 500 bp) were mapped to scaffolds (i.e., 99 ± 1.6% of reads mapped to scaffolds) (Table S3).

Open reading frames (ORFs) on scaffolds were predicted using Prodigal (Hyatt *et al*., 2010) with the meta flag activated. The resulting predicted ORFs were annotated against KEGG (Kyoto encyclopaedia of genes and genomes) (Kanehisa *et al*., 2015) using DIAMOND (Buchfink *et al*., 2015). Genes involved in the nitrogen cycle were identified based on KEGG orthology (KO) numbers assigned to predict ORFs based on the KEGG nitrogen metabolism pathway (Table S4). The abundance (as reads per million kilobase (rpkm)) of genes was determined across all samples by dividing the number of reads mapping to scaffold containing the gene by the scaling factor (i.e., millions of reads per sample) and the length of the scaffold in kilobases. The quality filtered paired end reads were mapped to a database of 44 high quality complete and draft genomes of nitrifying organisms, AOA (n=11), AOB (n=19), comammox bacteria (n=4), NOB (n=5), and anammox bacteria (n=5) (Table S5). Reads were competitively mapped to reference genomes using bwa and properly paired reads (samtools view –f 2) mapping to each reference were counted using awk script. The abundance of reference genomes in the samples was calculated by dividing the number of reads mapping to reference genomes by the scaling factor (i.e., millions of reads per sample) and the total length of the reference genome in kilobases. All raw sequence data have been deposited with links to BioProject accession number PRJNA524999 in the NCBI BioProject database (https://www.ncbi.nlm.nih.gov/bioproject/).

### 2.4 Metagenome Assembled Genome (MAG) reconstruction

Assembled scaffolds (>2000 bp) from the co-assembly of all samples were used to generate metagenomic assembled genomes (MAGs) using CONCOCT (Alneberg *et al*., 2014). This resulted in the construction of 115 CONCOCT clusters. A total of 60 CONCOCT clusters with completeness greater than 50%, based on the occurrence of 36 single copy genes used by CONCOCT to estimate completeness, were selected for further examination/refinement. The completeness and redundancy of these 60 CONCOCT clusters was checked with CheckM (Parks *et al*., 2015) with 47 clusters selected for further analysis based on 75% completeness. Of these, three had redundancy estimates greater than 10% and were manually refined using Anvi’o (Eren *et al*., 2015). This resulted in 47 high quality Metagenome Assembled Genomes (MAGs) (>70% complete, less than 10% redundancy). Bins were functionally annotated and taxonomically classified using GhostKOALA (Kanehisa *et al*., 2016), where predicted ORF’s were assigned KO numbers using KEGG’s set of nonredundant KEGG genes. In addition, the abundance of all MAGs was calculated similar to that of the reference genomes (i.e., rpkm). Taxonomic annotation of the final MAGs was conducted using MiGA (Rodriguez and Konstantinidis, 2014). Taxonomic inference and characteristics of MAGs are detailed in Table S6. Genome-level inference of the 47 MAGs was then used to construct a phylogenomic tree using GToTree described by Lee (2019).

### 2.5 Marker gene based taxonomic and phylogenetic analysis

Small subunit (SSU) rRNA gene sequences were identified using a Hidden Markov Model (HMM) search using the Infernal package (Nawrocki *et al*., 2009) with domain specific covariance models and corrected as outlined previously (Brown *et al*., 2015). Detected SSU rRNA genes greater than 500 bp were classified using SILVA taxonomy and the relative abundance of each SSU rRNA gene was estimated by dividing the total coverage of the scaffold containing the SSU rRNA gene by the coverage of all scaffolds containing SSU rRNA genes within each domain (i.e., bacteria, archaea, and eukaryota). Reference databases of ammonia moonooxygenase subunit A (*amoA*, KO: K10944) and nitrite oxidoreductase subunit A (*nxrA*, KO: K00370) genes were created using corresponding reference sequences obtained from NCBI GenBank Database with additional *nxrA* reference sequences obtained from Kitzinger *et al*. (2018). An alignment was created for each gene using MAFFT (version 7) online multiple alignment tool with the iterative refinement method L-INS-i (Katoh *et al*., 2002). Resulting alignments were examined and trimmed by removing all overhangs using BioEdit Sequence Alignment Editor v 7.2.6.1 (Hall, 2011), resulting in sequences of equal length. Aligned datasets were subjected to Maximum Likelihood analysis (Felsenstein, 1981) in MEGA7 (Molecular Evolutionary Genetic Analysis) (Kumar *et al*., 2016) using the best-fit substitution models as determined in MEGA7 model tests. For all Maximum Likelihood phylogenetic trees, branch support was estimated using non-parametric bootstrap analyses based on 1000 pseudoreplicates under the same model parameters and rooted with appropriate outgroups. Amino acid sequences of genes annotated as *amoA* and *nxrA* were then placed on the respective reference phylogenetic tree using pplacer (Matsen *et al*., 2010).

## 3. Results

### 3.1 Changes in ammonium, nitrite, and nitrate concentrations in the chloraminated section of the DWDS

Spatial changes in the concentrations of ammonium, nitrite and nitrate were observed as the chloraminated bulk water moved through the DWDS (Fig. S1). Here, ammonium concentrations decreased as the bulk water moved away from the chloramination sites towards the end of the DWDS (i.e., average NH_4_^+^ concentrations decreased from 0.30 ± 0.12 mg/L following chloramination to 0.05 ± 0.09 mg/L at the end of the DWDS). The decrease in ammonium concentrations was associated with an increase in nitrite concentrations, which reached its highest concentrations at sites before RES2 (i.e., average NO_2_^−^ concentrations increased from 0.005 ± 0.01 mg/L to 0.21 ± .033 mg/L) after which they decreased. Interestingly, both nitrite and nitrate levels increased in RES1, while increases in nitrate concentrations directly corresponded to decreases in nitrite concentrations after RES2. Nitrate concentrations peaked at locations after RES2 where ammonium and nitrite concentrations are the lowest (i.e., average NO_3_^−^ concentrations increased from 0.18 ± 0.12 mg/L following chloramination to 0.40 ± 0.23 mg/L at the end of the DWDS).

A clear decrease in ammonium concentrations was observed after RES2 (Fig. 2B) (i.e., average ammonium concentrations of 0.30 ± 0.13 mg/L before RES2 and 0.16 ± 0.12 mg/L after RES2). In the case of RES1, increases in nitrite and nitrate concentrations were generally associated with concomitant decreases in ammonium concentrations in the first 8 months. However, these correlations between nitrite, nitrate and ammonium concentrations were not observed for the remaining months of the study period. Samples before RES2 had increased nitrite and nitrate concentrations and lower ammonium concentrations as compared to RES1. However, after RES2 nitrite levels dramatically decrease (i.e., average NO_2_^−^ concentrations decreased from 0.17 ± 0.22 mg/L before RES2 to 0.09 ± 0.09 mg/L after RES2), which correlated with an increase in nitrate concentrations (i.e., average NO_3_^−^ concentrations increased from 0.27 ± 0.13 mg/L before RES2 to 0.41 ± 0.22 mg/L after RES2).

**Fig. 2:**
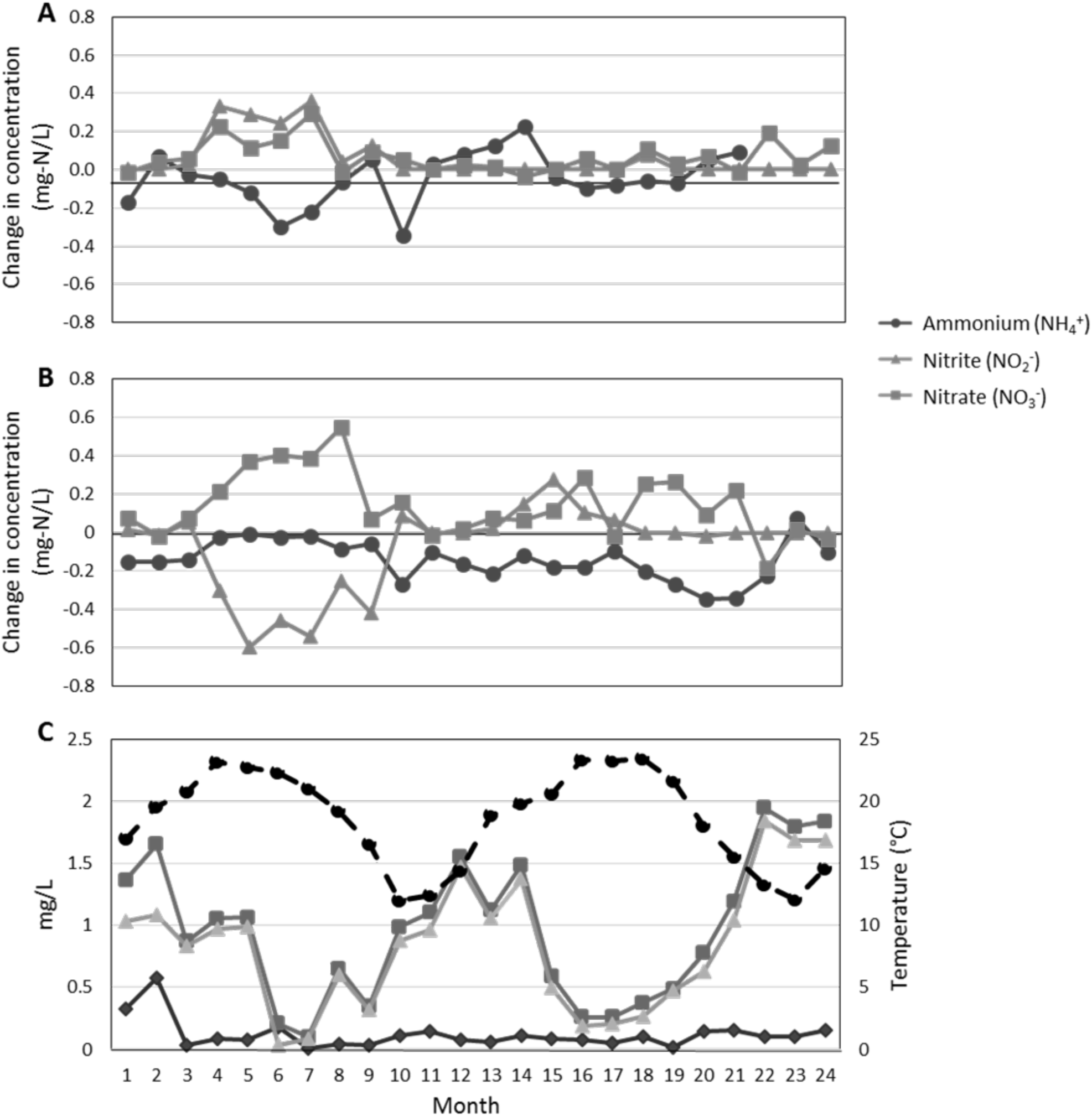
Change in nitrogen species concentration [i.e., ammonium (red circles), nitrite (green triangles) and nitrate (blue squares)] (A) before and after reservoir 1 (RES1) and (B) before and after reservoir 2 (RES2) over the two year sampling period. Average concentrations of disinfectant residuals [i.e., free chlorine (diamonds), total chlorine (squares) and monochloramine (triangles)] together with average temperature (dashed black line with circles) across RES1 and RES2 (C).

Furthermore, ammonium, nitrite and nitrate concentrations demonstrated strong temporal trends in both reservoirs associated with changes in temperature and disinfectant residual concentrations with the highest ammonium concentrations in winter and spring months (~0.5 mg/l). Disinfection residual concentrations (i.e. total chlorine and monochloramine) were generally higher in winter and spring months (peaking in July 2016 at approximately 2.0 mg/L) and negative correlated with water temperature as shown previously (Potgieter *et al*., 2018) (Fig. 2C). Nitrite and nitrate concentrations were highest in summer and autumn months (0.66 and 0.82 mg/L, respectively) where associated ammonium levels were low (0.12 mg/L). Typically, decrease in monochloramine and ammonium concentrations were associated with increased nitrite and/or nitrate concentrations at higher temperatures. It is important to note that the observed trends in ammonium, nitrite and nitrate concentrations may not be the same from year to year as increased levels of nitrification were observed for the first year (months 10-12) as opposed to second year (months 13-24) (Fig. S2).

### 3.2 Changes in the microbial community composition between the two reservoirs

Based on SSU rRNAs identified in the metagenomic data, the majority of sequences identified were bacterial (i.e., the mean relative abundance (MRA) of bacterial SSU rRNA genes across all samples was 90.72 ± 7.23%), followed by unclassified SSU rRNA contigs (6.58 ± 5.29%) and eukaryota (2.70 ± 2.65%) with no archaeal SSU rRNA detected (Table S7). At the phylum level, *Proteobacteria* were the most abundant in both reservoirs (i.e., MRA of 81.60 ± 13.32% in RES1 and 82.98 ± 10.28% in RES2), followed by *Nitrospirae* with an MRA of 11.25 ± 13.48% in RES1, although the MRA of *Nitrospirae* decreased to 2.28 ± 3.83% in RES2 (Fig. S3A). The decrease in abundance of both *Proteobacteria* and *Nitrospirae* in RES2 was associated with increases in both eukaryota (MRA: 2.05 ± 1.26% in RES1 to 4.64 ± 5.55% in RES2) and unclassified contigs (MRA: 3.55 ± 1.17% in RES1 to 7.58 ± 3.66% in RES2), of which 16.87% of unclassified contigs were less than 250 bp. The taxonomic classification of eukaryota 18S rRNA contigs is shown in Table S8.

Further classification of the proteobacterial classes based on SILVA taxonomy revealed that the abundance of *Alpha*- and *Gammaproteobacteria* (specifically order: *Betaproteobacteriales*) varied between RES1 and RES2. In RES1 the abundance of *Alphaproteobacteria* (MRA: 42.99 ± 13.97%) was marginally higher than *Betaproteobacteriales* (MRA: 38.05 ± 14.47%). However in RES2, the abundance of *Alphaproteobacteria* increased in dominance (MRA: 52.75 ± 15.36%) and *Betaproteobacteriales* decreased (MRA: 29.46 ± 17.14%) indicating a decrease in the abundance of *Betaproteobacteriales* as chloraminated water moves further down the DWDS.

In RES2 there was an observed increase in the relative abundance of other *Gammaproteobacteria* from a MRA of 0.48 ± 0.37% in RES1 to 0.69 ± 0.35% in RES2 (Fig. S3B). Furthermore, within *Betaproteobacteriales*, two *Nitrosomonas* SSU rRNAs (NODE_284 and NODE_310) were highly abundant with the mean relative abundance of 31.44 ± 15.18% in RES1 and 10.10 ± 10.07% in RES2, consequently making *Nitrosomonas* the most dominant genera identified in all samples. These results correlated with 16S rRNA gene profiling data previously reported by Potgieter *et al*. (2018).

Alpha diversity indices [richness (observed taxa), Shannon Diversity Index and Pielou’s evenness] were calculated using the summary single function in mothur (Schloss et al., 2009) incorporating the parameters, iters=1000 and subsampling=475 (sample containing the least number of sequences). RES2 samples were more rich than RES1 samples (average number of taxa for RES1 was 49 ± 12, whereas for RES2 it was 69 ± 28) as well as slightly more diverse and even (i.e., Shannon Diversity Index: 2.91 ± 0.77 and Pielou’s evenness: 0.69 ± 0.13) than RES1 (i.e., Shannon Diversity Index: 2.32± 0.44 and Pielou’s evenness: 0.59 ± 0.10) (Fig. S4). However, Kruskal-Wallis one-way analysis of variance and post-hoc Dunn’s test revealed that differences in alpha diversity measures between the two reservoirs were not significant (p > 0.05). Beta diversity measures (i.e. structure based: Bray-Curtis and membership based: Jaccard) between the two reservoirs for corresponding months revealed that the microbial communities between the two reservoirs were on average 60% dissimilar in community structure (Bray-Curtis: 0.59 ± 0.16) and 74% dissimilar in community membership (Jaccard: 0.74 ± 0.12). This indicated that the microbial community differs significantly in both community structure and membership between the two reservoirs (AMOVA, *F_ST_* ≤ 2.12, p < 0.05).

### 3.3 Dominant genes involved in nitrogen transforming reactions

Genes encoding for enzymes involved multiple nitrogen transforming reactions were observed across both reservoirs (Table S4). However, the coverage of some of these genes was low and as a result their contribution to overall nitrogen metabolism was thought to be limited in the chloraminated drinking water environment. Therefore, the dominant genes driving nitrogen transformation reactions were identified as those genes with a cumulative coverage of >100 reads per million kilobase (rpkm) in both reservoirs (Fig. 3). These genes included *amoABC* and *hao* (ammonia oxidation), *nxrAB* (nitrite oxidation), *nasA* (assimilatory nitrate reduction), *nirBD* (assimilatory nitrite reduction), *nirK* (nitrite reduction, NO-forming) and *norBCDQ* genes (nitric oxide reduction). The cumulative coverage of these dominant genes for each sample from both reservoirs in shown in Fig. 4.

**Fig. 3:**
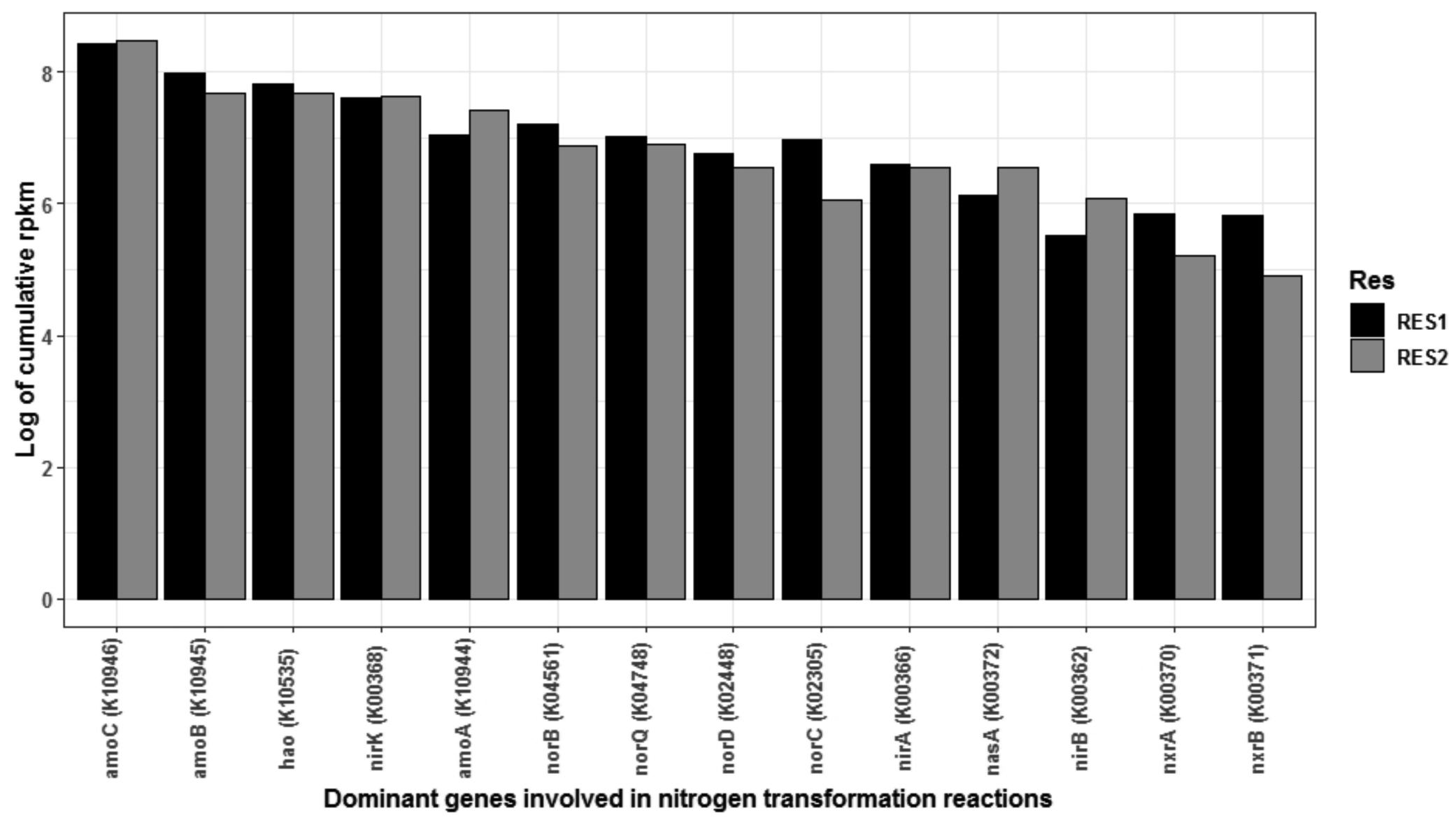
Log transformed cumulative coverage (rpkm) of the dominant genes identified to be involved in the nitrogen cycle (i.e. genes with a cumulative coverage of > 100 rpkm in both reservoirs).

**Fig. 4:**
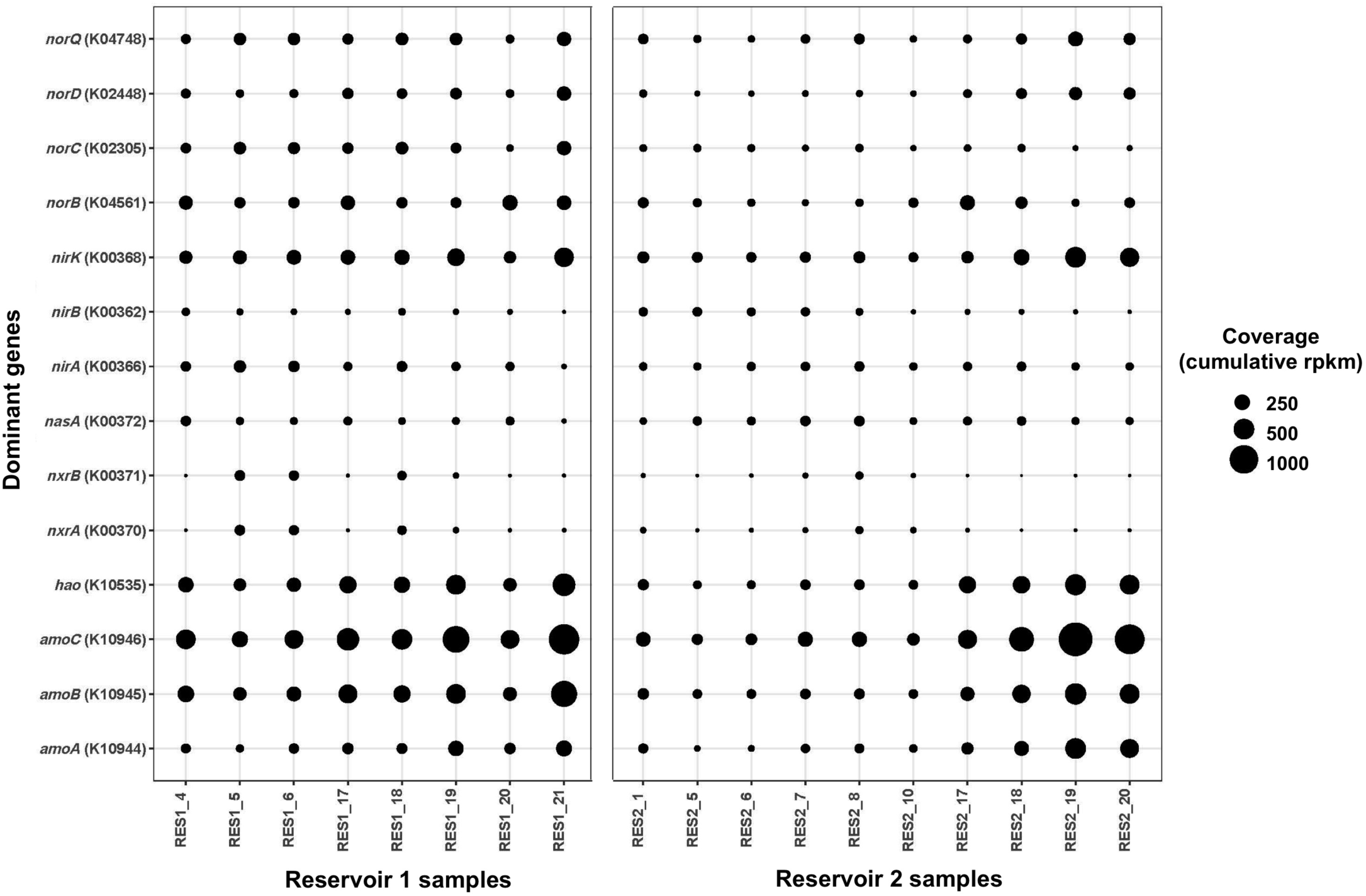
Cumulative coverage (rpkm) of the dominant genes identified to be involved in the nitrogen cycle (i.e. genes with a cumulative coverage of >100 rpkm in both reservoirs) across all reservoir samples. This figure is complemented by Fig. S5, showing the number of genes identified for each function.

#### 3.3.1 Ammonia and nitrite oxidation

Genes encoding for the enzymes involved in nitrification were observed in all reservoir samples. Specifically, the genes responsible for ammonia oxidation were observed to have the highest coverage across all samples, out of all genes identified to be involved in the nitrogen cycle (Fig. 4). In both reservoirs, *amoC*, *amoB* and *hao* genes had the highest total cumulative coverage in RES1 (i.e. 4625, 2922 and 2486 rpkm, respectively) and in RES2 (i.e. 4807, 2166 and 2170 rpkm, respectively). The average cumulative coverage of these dominant genes was higher across RES1 samples (i.e. *amoC*: 578 ± 287, *amoB*: 365 ± 206 rpkm and *hao*: 310 ± 150 rpkm) than across RES2 samples (i.e. *amoC*: 481 ± 458, *amoB*: 217 ± 171 rpkm and *hao*: 217 ± 172 rpkm). In addition, *amoA* genes were also observed to be highly abundant with a total cumulative coverage of 1143 rpkm in RES1 for all time points (average cumulative coverage of 143 ± 80 rpkm across all RES1 samples), which increased to 1672 rpkm in RES2 (with an average cumulative coverage of 167 ± 163 rpkm across all RES2 samples) (Fig. 3 and 4). Although, these genes showed high coverage, only a small number of genes were identified as *amoABC* and *hao*, indicating a low level of diversity among these genes (Fig. S5). Here, the four individual contigs with the highest coverage across all samples (i.e. average coverage across all samples > 100 rpkm) contained the *hao* (NODE_10017), *amoC* (NODE_69075), *amoB* (NODE_58840) and *amoABC* (NODE_21034) genes. BLAST results revealed that the majority of *amoABC* and *hao* genes represented members of *Nitrosomonas* genus with a small minority of genes within each group were found to belong to *Nitrospira* species, indicating the potential presence of comammox bacteria in the bacterial community.

Genes encoding nitrite oxidoreductase (*nxrAB*), required for the oxidation of nitrite to nitrate, were also observed across all samples (Fig. 4). As expected, BLAST results of *nxrA* and *nxrB* genes revealed that the majority of these genes represented *Nitrospira* species. Although the coverage of individual *nxrAB* genes was significantly lower than *amoABC* and *hao* genes, they still maintained a cumulative coverage of >100 rpkm in both reservoirs. *nxrA* genes showed a total cumulative coverage of 347 rpkm in RES1 (average cumulative coverage of 43 ± 45 rpkm across RES1 samples) and decreased to 183 rpkm in RES2 (with an average cumulative coverage of 18 ± 16 rpkm across RES2 samples). Similarly, *nxrB* genes showed a total cumulative rpkm of 335 rpkm in RES1 (average cumulative coverage of 42 ± 46 rpkm across RES1 samples) and decreased to 135 rpkm in RES2 (with an average cumulative coverage of 14 ± 17 rpkm across RES2 samples) (Fig. 3 and 4). Specifically, the individual contig NODE_68 containing *nxrAB* genes was consistently present across all samples and contributed to the high cumulative coverage of *nxrAB* genes (with an average cumulative coverage of 39 ± 48 rpkm in RES1 and 9 ± 16 rpkm in RES2). The low coverage of *nxrAB* genes correlated with the small number of genes identified as *nxrAB* genes, indicating a low level of diversity within this function (Fig. S5).

Although different in each reservoir, the temporal trends in the cumulative coverage revealed a contrasting relationship between *amoABC* and *nxrAB* genes (Fig. S6A and S6B). In RES1, *nxrAB* genes showed increased coverage in March (2015 and 2016) relative to *amoA* genes. Conversely, months where *nxrAB* gene coverage decreased, the coverage of *amoABC* genes increased (April 2015 and 2016). This contrasting temporal trend was also observed in RES2.

#### 3.3.2 Reduction of nitrate and nitrite

The nitrate formed through nitrification can be potentially reduced back to nitrite through nitrate reduction. Here, the cytoplasmic assimilatory nitrate reductase, *nasA* gene (encoding the catalytic subunit of the NADH-nitrate reductase) was identified as the most abundant nitrate reductase, indicating the capacity of certain members of the community to use nitrate as an alternative electron acceptor. Dissimilatory nitrate reductase genes were also identified, including respiratory membrane bound nitrate reductases (*narGHIJ*), although their cumulative coverage of was low (<100 cumulative rpkm in both reservoirs). The total cumulative coverage of all *nasA* genes identified was 455 rpkm in RES1 (average cumulative coverage of 57 ± 24 rpkm across RES1 samples) and increased to 701 rpkm in RES2 (with an average cumulative coverage of 70 ± 22 rpkm across RES2 samples) (Fig. 3 and 4). BLAST analyses revealed that the majority of *nasA* genes represented members of *Alphaproteobacteria* (6% of *nasA* genes) of which 34% were identified as belonging to the order *Rhizobiales*. In addition, *Gammaproteobacteria* (order: *Betaproteobacteriales*) represented another 18% of *nasA* genes. Here, an increased number of genes within the metagenomic dataset were identified as *nasA*, converse to *amo* and *nxr* genes, indicating an increased level of diversity of microorganisms containing the cytoplasmic assimilatory nitrate reductase (Fig. S5).

Genes encoding enzymes involved the assimilatory reduction of nitrite to ammonia (*nirAB* genes) were also observed to have high coverage across all reservoir samples (Fig. 4). The ferredoxin-nitrite reductase (*nirA*) showed high abundance with a total cumulative coverage of 729 rpkm in RES1 (average cumulative coverage of 91 ± 41 rpkm across RES1 samples) and 629 rpkm in RES2 (with an average cumulative coverage of 69 ± 13 rpkm across RES2 samples) (Fig. 3 and 4). BLAST analyses identified that 50% of *nirA* genes represented members of *Alphaproteobacteria* of which 36% were identified as *Rhizobiales* while 32% of *nirA* genes were identified as belonging to *Nitrospira* species. The nitrate reductase (NADH), large subunit (*nirB* gene) was also observed to have high coverage across both reservoirs (i.e. cumulative coverage >100 rpkm), although the abundance of *nirB* genes was less than that of the *nirA* genes with the total cumulative coverage of *nirB* genes being 249 rpkm in RES1 (average cumulative coverage of 32 ± 16 rpkm across RES1 samples) and increased to 438 rpkm in RES2 (with an average cumulative coverage of 44 ± 31 rpkm across RES2 samples) (Fig. 4). BLAST results revealed that the majority of *nirB* genes represented *Gammaproteobacteria* (order: *Betaproteobacteriales*) (69% of *nirB* genes) of which 28% were identified as *Nitrosomonadales* and 28% as *Burkholderiales*. In addition, another 21% of *nirB* genes were found to represent *Alphaproteobacteria* (order: *Rhizobiales*). Although, both *nirAB* genes showed moderate to low coverage across both reservoirs, the number of genes identified as belonging to these functions was high, specifically *nirB* genes. This indicates that a diverse assemblage of bacteria had the genetic potential for assimilatory nitrite reduction (Fig. S5).

However in RES1, temporal trends in the relative abundance of *nasA* and *nirA* genes showed a converse relationship (Fig. S6C). Increased relative abundance of *nirA* genes (in February and March 2015 and March 2016) was associated with decreased relative abundance of *nasA* genes. However, this converse relationship was not observed between *nasA* and *nirB* in RES2. In RES2, the relative abundance of these genes (*nasA*, *nirA* and *nirB*) generally showed the same temporal trends across RES2 samples and the same converse relationship with ammonia concentrations, specifically between *nasA* and *nirA*. Increases in the relative abundance of both *nasA* and *nirA* genes was observed in April and May 2015 and March 2016 (Fig. S6C). This suggests the potential for complete assimilatory nitrate reduction to ammonia (i.e., *nasA* and *nirAB*) specifically in RES2 as the relative abundance of *nasA* genes typically showed a converse relationship to ammonia concentrations.

Lastly, *nirK* genes encoding nitric oxide forming nitrite reductases (reducing nitrite to nitric oxide) were also observed to be highly abundant in both reservoirs (i.e. cumulative coverage >100 rpkm) (Fig. 3). *nirK* genes were identified as the fourth most abundant gene (following *amoC*, *amoB* and *hao*, respectively) with total cumulative coverage of 2016 rpkm in RES1 (with an average cumulative coverage of 252 ± 92 rpkm across RES1 samples) and 2043 rpkm in RES2 (with an average cumulative coverage of 204 ± 114 rpkm across RES2 samples) (Fig. 3 and 4). Furthermore, the cumulative coverage of *nirK* genes across all samples generally showed a converse relationship with *nirA* and *nirB* genes in both reservoirs (Fig. S6C and S6D). In addition, a high number of genes were identified as *nirK*, indicating a high diversity within this function (Fig. S5). Furthermore, BLAST results of individual *nirK* genes revealed that 36% of these genes belonged to species of *Nitrospira* and 17% to *Nitrosomonas* species. The remaining *nirK* genes were shown to represent a diverse group of bacteria including both *Alpha*- and *Gammaproteobacteria* (order: *Betaproteobacteriales*).

#### 3.3.3 Nitric oxide reduction

The nitric oxide formed through the reduction of nitrite can be further reduced to nitrous oxide by nitric oxide reductases (*norBCDEQ*). Genes encoding for nitric oxide reductases (*norBCDQ*) were identified to have high coverage in this dataset, across all samples (i.e. cumulative coverage >100 rpkm in both reservoirs) (Fig. 3). These genes had higher total cumulative coverage in RES1 (i.e. *norB*: 1364 rpkm, *norQ*: 1122 rpkm, *norC*: 1060 rpkm and *norD*: 856 rpkm) than in RES2 (i.e. *norB*: 972 rpkm, *norQ*: 997 rpkm, *norC*: 427 rpkm and *norD*: 700 rpkm) (Fig. 3 and 4). Interestingly, the temporal cumulative coverage of *norB* generally showed a converse relationship to the cumulative coverage of other *nor* genes, specifically *norC* and *norQ* (Fig. S6E). The dynamics of *nor* genes may be associated with different members of the community therefore resulting in differences in their coverage. The diversity of *norBCQ* genes was generally higher than that of *norD* genes (Fig. S5). BLAST analyses revealed that *norB* genes represented members of *Gammaproteobacteria* (order: *Betaproteobacteriales*) predominately the order *Nitrosomonadales* (33% of *norB* genes) and *Alphaproteobacteria*, predominately the family *Sphingomonaceae* (23% of *norB* genes). The majority of *norQ* genes were identified as *Nitrospira* (16% of *norQ* genes) and *Gammaproteobacteria* (order: *Betaproteobacteriales*) predominately the order *Nitrosomonadales* (40% of *norQ* genes), of which 14% was identified as *Nitrosomonas* species. Similarly, the majority of *norC* genes were identified as *Nitrospira* (44% of *norC* genes) and *Gammaproteobacteria* (order: *Betaproteobacteriales*) order *Nitrosomonadales* (36% of *norC* genes). Lastly, as with *norQ* and *norC* genes, 55% *norD* genes were found to represent members of *Gammaproteobacteria* (order: *Betaproteobacteriales*) order *Nitrosomonadales* (of which 21% was identified as *Nitrosomonas* species) and 13% as members of the genus *Nitrospira*.

#### 3.3.4 Other processes involved in nitrogen metabolism

Briefly, genes involved in other processes in nitrogen metabolism, with very low coverage across both reservoirs (< 100 cumulative rpkm) also were identified (Table S4). These genes included nitrous oxide reductase (*nosZ*), nitrogenases involved in nitrogen fixation (*nif* genes) as well as other genes involved in nitrite and nitrate transport (i.e. *nrtABC* genes encoding the enzymes periplasmic substrate-binding protein, permease protein and an ATP-binding protein, respectively). The presence of *nrtABC* often co-occurred with presence of *nasA*, suggesting these nitrate transport enzymes are associated with assimilatory nitrate reductases. In addition, the gene encoding nitronate monooxygenase (*ncd*2) was also identified suggesting the potential to convert nitronate to nitrite and potentially provide an alternative source of nitrite.

### 3.4 Nitrifier diversity based on phylogenetic inference of *amoA* and *nxrA* genes

Given the high abundance of *Nitrosomonas* and *Nitrospira*, the phylogenetic diversity of *amoA* and *nxrA* genes (marker genes for ammonia and nitrite oxidation, respectively) was investigated. Consistent with BLAST results, the phylogenetic placement of *amoA* genes revealed that nearly all *amoA* genes clustered with *Nitrosomonas* species indicating that the majority of *amoA* genes are associated with strict ammonia oxidisers (Fig. 5A). Of these genes, three grouped closely with *Nitrosomonas oligotropha*, one with *Nitrosomonas* sp. AL212 and one with *Nitrosomonas* sp. Is79A3. The clustering of the remaining contigs remained within the *Nitrosomonas* but did not group closely with specific species (Fig. 5A). Interestingly, a single *amoA* contig (NODE_6011) grouped within the *Nitrospira* comammox cluster. Furthermore, BLAST results confirmed that *amoA* contig (NODE_6011) represented a *Nitrospira* species, indicating the potential presence of a comammox bacteria within the microbial community.

**Fig. 5:**
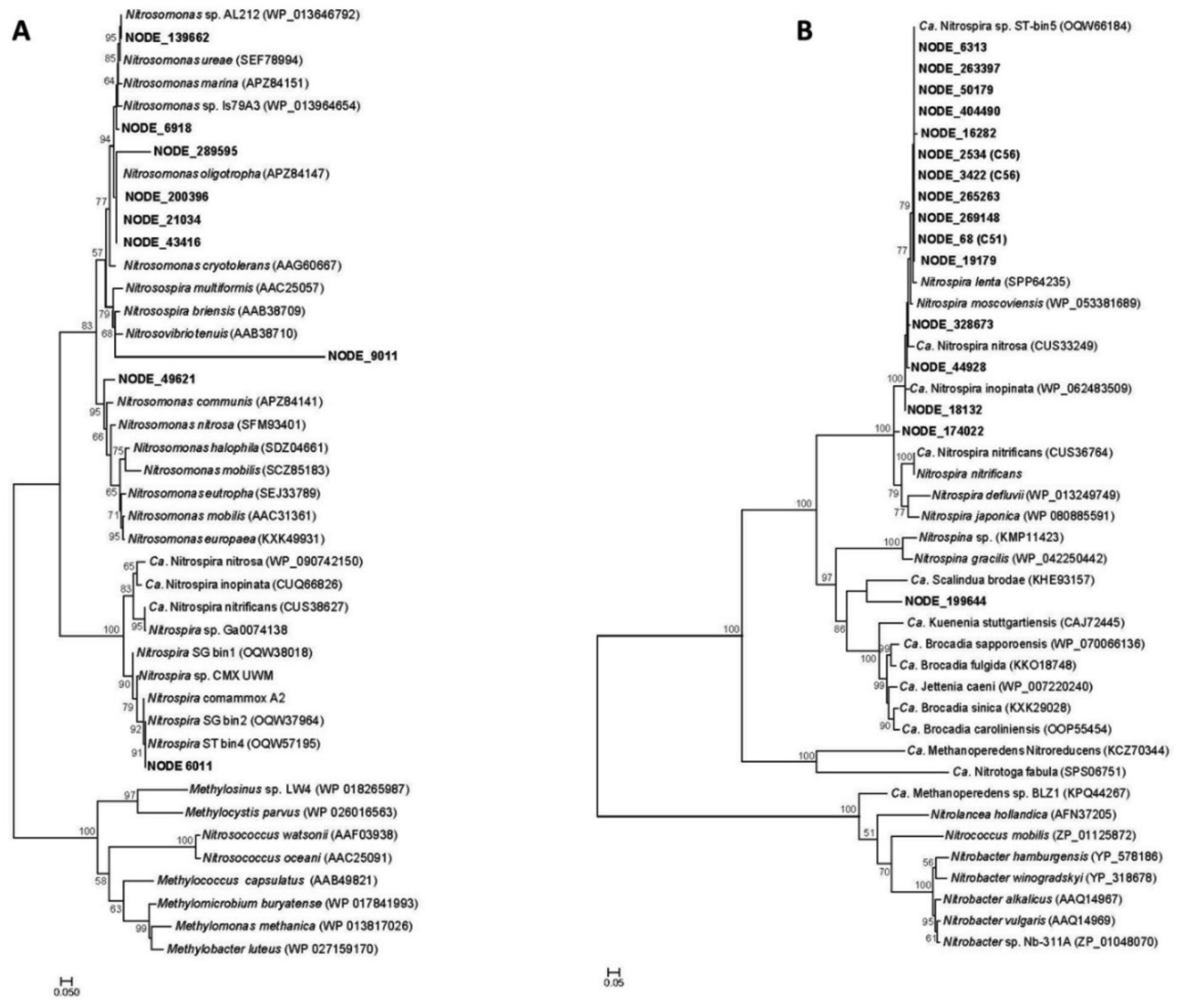
Phylogenetic placement of (A) ammonia monooxygenase, subunit A (*amoA*) and (B) nitrite oxidoreductase, subunit A (*nxrA*) phylogenies. Both Maximum Likelihood phylogenetic trees were constructed based on amino acid sequences from contigs identified as the respective genes. Contigs identified from this study are in bold. Reference trees were constructed with bootstrap analysis of 1000 replicates. Bootstrap values are indicated as percentages and values below 50 were excluded.

Also consistent with BLAST results, phylogenetic placement of the *nxrA* genes revealed that the majority *of nxrA* genes belonged to the widely distributed lineage II of the genus *Nitrospira* grouping with the canonical nitrite oxidisers, *N. lenta* and *N. moscoviensis* and *Ca*. Nitrospira sp. ST-bin5 described by Wang *et al*. (2017) (Fig. 5B). In addition, some *nxrA* genes grouped closely with known comammox *Nitrospira*, including *Ca*. Nitrospira nitrosa, *Ca*. Nitrospira inopinata and *Ca*. Nitrospira nitrificans. However, identification and taxonomic resolution of comammox bacteria based on *nxrA* gene phylogeny is not possible. Lastly, a single *nxrA* gene grouped closely with the anammox *Ca*. Scalindua brodae although no other evidence of anammox bacteria was found in the metagenomic data. Furthermore, no *nxrA* genes were found to be associated with *Nitrobacter* species (Fig. 5B).

### 3.5 The nitrogen metabolic potential of dominant Metagenome Assembled Genomes (MAGs)

Following metagenomic binning, the 47 high quality MAGs constructed included 25 *Alphaproteobacteria*, 15 *Betaproteobacteriales*, 2 *Nitrospirae* as well as another 5 MAGs identified as *Bacteroidetes*, *Gammaproteobacteria*, *Gemmatimonadetes* and 2 *Planctomycetes*, respectively (Fig. 6). MAGs were genetically compared to their closest related reference genomes based on average amino acid identity (AAI). The complete description of the 47 MAGs and their coverage across the two reservoirs are shown Fig. S7 and described in Table S6. This analysis suggested that the majority of MAGs constructed represent genomes that are not yet represented in public database. However, the classification of these MAGs identified members of several abundant taxa identified in this chloraminated DWDS described by Potgieter *et al*. (2018). The 47 constructed draft genomes were screened for key functional genes involved in nitrogen metabolism described in Table S4 and the genetic potential for nitrogen transforming reactions within these MAGs was found to be diverse (Fig. 6).

**Fig. 6:**
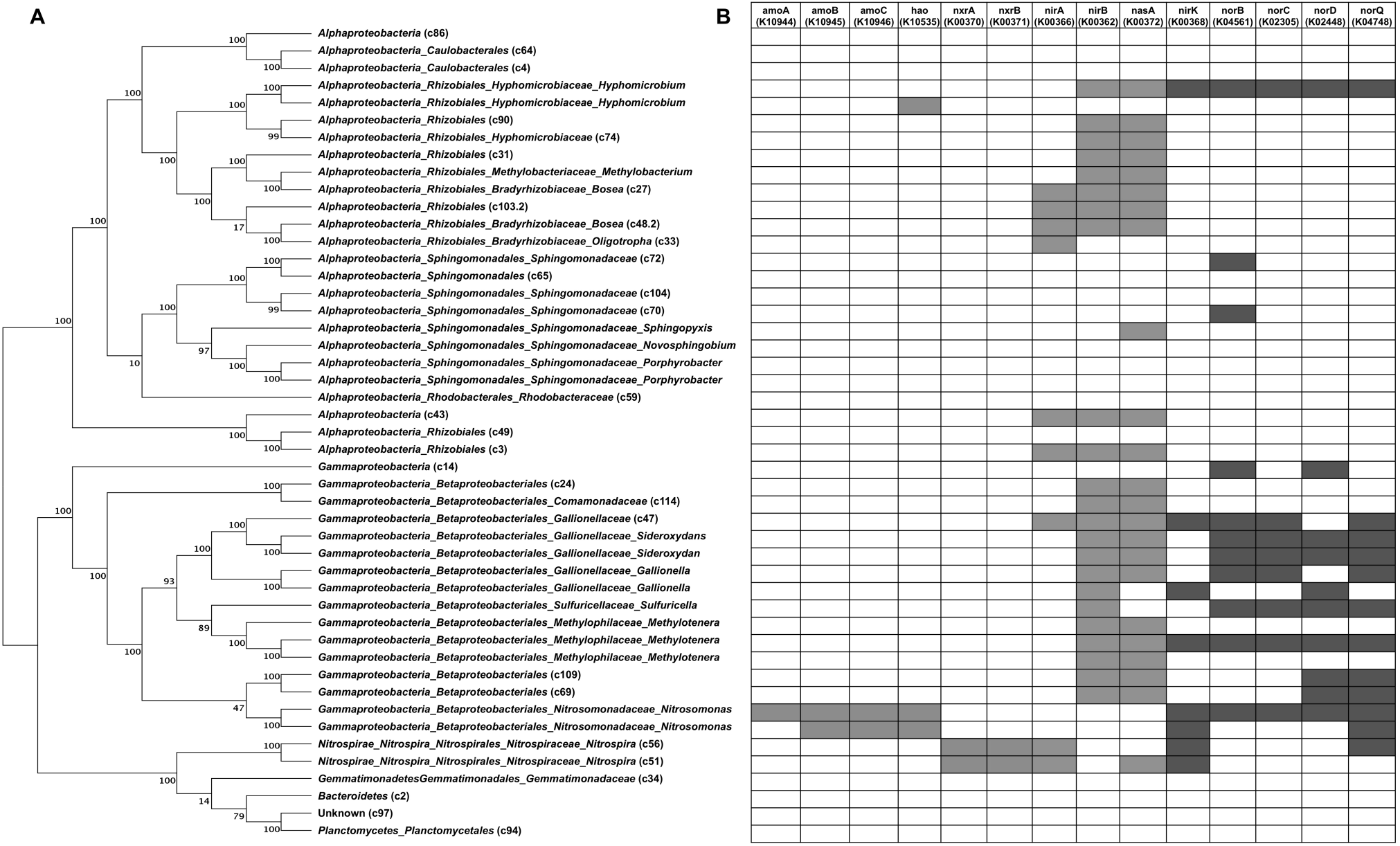
(A) Phylogenomic tree showing the genome-level evolutionary inference of the 47 MAGs constructed from the metagenomic dataset and (B) the corresponding presence of dominant genes associated with the nitrogen cycle identified in each MAG. (Green indicates those genes involved in nitrification, blue indicates genes involved in nitrate and nitrite reduction and red indicates genes involved in nitric oxide formation and reduction).

Of the 47 MAGs, 5 MAGs were observed to dominate the community with a cumulative rpkm of >100 rpkm across both reservoirs (Fig. 7). These MAGs were identified as two *Nitrosomonas*-like MAGs (C58 and C107), a *Rhizobiales*-like MAG (C103.2), a *Sphingomonas*-like MAG (C70) and a *Nitrospira*-like MAG (C51). Furthermore, the relative abundance of these top 5 most abundant SSU rRNAs in the metagenomic dataset were correlated with the rpkm based abundance of these MAGs showing the same trends across both reservoirs (Fig. S8) and the contigs containing these SSU rRNAs were observed in their corresponding MAGs.

**Fig. 7:**
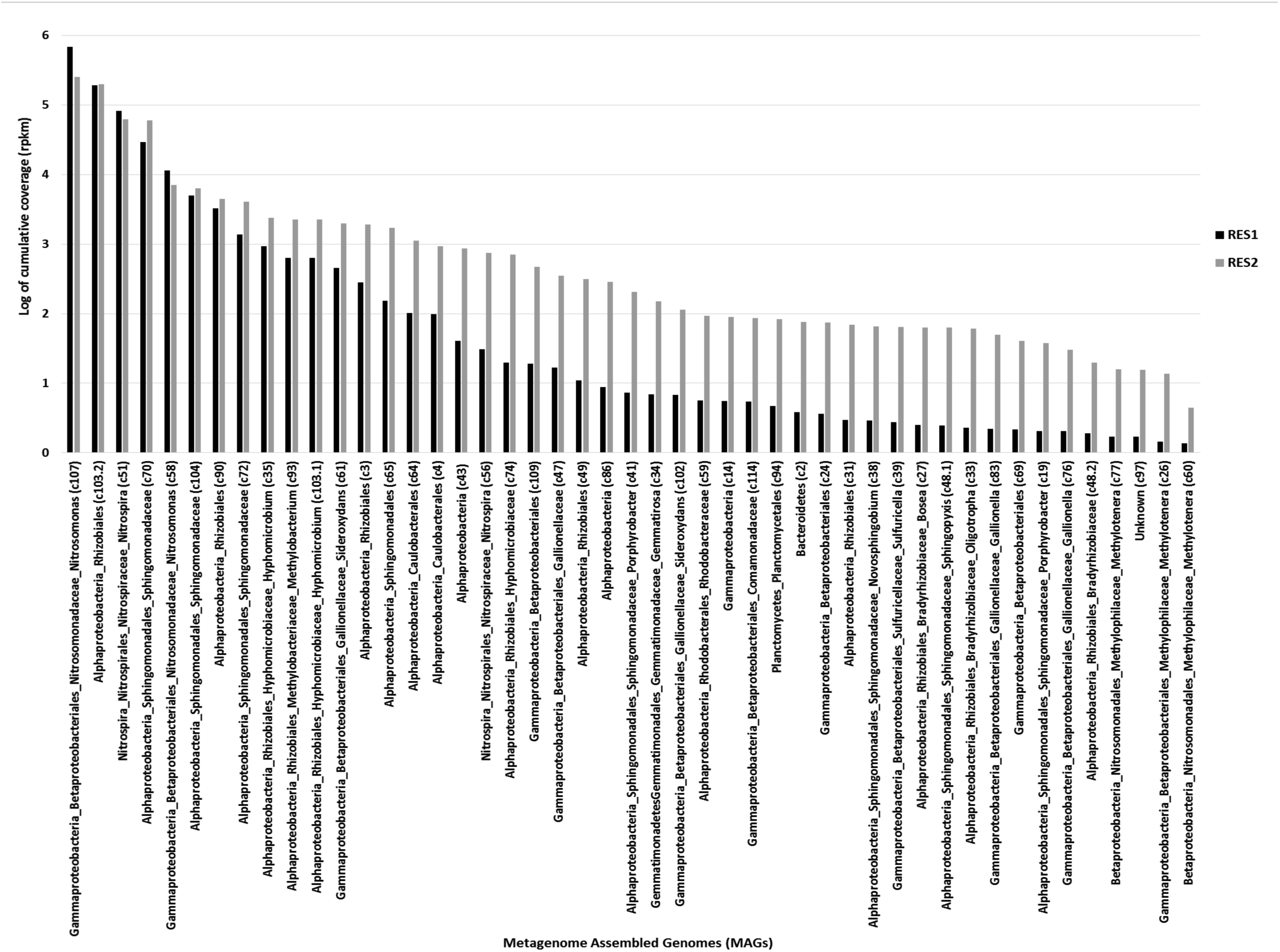
Log transformed cumulative coverage (rpkm) of the 47 reconstructed Metagenome Assembled Genomes (MAGs) for both reservoirs.

#### 3.5.1 *Nitrosomonas*-like MAGs (C58 and C107)

Contigs containing the *Nitrosomonas* SSU rRNAs were found to be associated with the two dominant *Nitrosomonas*-like MAGs [i.e. contig NODE_284 with the *Nitrosomonas*-like MAG (C58) and contig NODE_310 with the *Nitrosomonas*-like MAG (C107)]. Phylogenetic analysis, of these *Nitrosomonas* spp. SSU rRNA contigs revealed that they both clustered with *Nitrosomonas oligotropha*, with good bootstrap support (Fig. S9). The classification of the *Nitrosomonas* SSU rRNAs as *Nitrosomonas oligotropha* correlates with the high coverage observed of the *Nitrosomonas oligotropha* reference genome across all samples (i.e., average coverage of 47 ± 23 rpkm in RES1 and 29 ± 24 rpkm in RES2) (Fig. S11). Thus, there may be two separate populations of *Nitrosomonas* responsible for the first step in nitrification.

Of the dominant MAGs, *Nitrosomonas*-like MAG (C107) exhibited the highest abundance in the metagenomic dataset. Although the two *Nitrosomonas*-like MAGs were potentially identified as the same species (based on SSU rRNAs), differences in the average coverage of these MAGs across the two reservoirs indicated that they may exhibit competitive dynamics (Fig. S7 and S8). Here, the coverage of *Nitrosomonas*-like MAG (C58) increased from RES1 (7 ± 10 rpkm) to RES2 (22 ± 27 rpkm), whereas *Nitrosomonas*-like MAG (C107) decreased in coverage from RES1 (43 ± 26 rpkm) to RES2 (12 ± 6 rpkm) (Fig. 7). Spearman correlations revealed that there was a moderate negative correlation between these two MAGs in RES1, however this was not statistically significant (Spearman correlation: −0.40, p = 0.3268).

Within both the *Nitrosomonas*-like MAGs, genes required for ammonia oxidation, i.e., ammonia monooxygenase (*amoABC*) and hydroxylamine oxidoreductase (*hao*) were observed. More specifically, the dominant genes in the metagenomic dataset, *hao* (NODE_10017) and *amoABC* (NODE_21034) could be linked to the *Nitrosomonas*-like MAG (C107). Furthermore, the *amoA* contig (NODE_21034) was observed to group closely with *N. oligotropha* (Fig. 5A). However, an *amoA* gene was not recovered from *Nitrosomonas*-like MAG (C58) and this may be due to its overall lower abundance. In addition, both *Nitrosomonas*-like MAGs also contained *nirK* genes (nitrite reductase, nitric oxide forming) and *nor* genes (nitric oxide reductases) (Fig. 6). More specifically, *Nitrosomonas*-like MAGs (C107) contained multiple *nor* genes including *norBCDQ*, whereas *Nitrosomonas*-like MAGs (C58) only contained *norQ*, suggesting that these dominant MAGs may play a role in regulating the concentrations of nitric oxide.

Spearman correlations between *Nitrosomonas*-like MAG (C107) and other dominant MAGs revealed strong correlations specifically in RES2. Strong positive correlations were observed with *Sphingomonas*-like MAG (C70) (Spearman correlation: 0.76, p < 0.05) and *Rhizobiales*-like MAG (C103.2) (Spearman correlation: 0.72, p < 0.05). In addition, *Nitrosomonas*-like MAG (C58) showed a strong positive correlation with *Rhizobiales*-like MAG (C103.2) (Spearman correlation: 0.81, p < 0.05) (Fig. S8).

#### 3.5.2 *Nitrospira*-like MAG (C51)

The *Nitrospira*-like MAG (C51) was dominant across all samples showing the 5th highest average coverage (10 ± 15 rpkm). The coverage of *Nitrospira*-like MAG (C51) was higher across RES1 samples (17 ± 20 rpkm) but decreased in RES2 (4 ± 69 rpkm) (Fig. 7). This same trend was observed with the *Nitrospira* SSU rRNA contig (NODE_2) as the contig was identified in *Nitrospira*-like MAG (C51). Following phylogenetic analysis of *Nitrospira* SSU rRNAs, it was observed that *Nitrospira* spp. SSU rRNA NODE_2 grouped closely with *Nitrospira lenta* (lineage II) (Fig. S10). The classification of the dominant *Nitrospira* SSU rRNA as *Nitrospira lenta* correlated with the observation of high coverage of the *Nitrospira lenta* reference genome across all samples with an average coverage of 14 ± 15 rpkm in RES1 and 4 ± 5 rpkm in RES2 (Fig. S11). Although the SSU rRNA (NODE_2) was identified as *Nitrospira lenta* and this *Nitrospira*-like MAG (C51) does not contain *amoABC* or *hao* genes, further investigation into the potential that this MAG was a comammox revealed that it does contain genes for cytochrome *c* biogenesis, *ccmEFGH* genes, thought to be specific to comammox *Nitrospira*. However, BLAST results revealed that these *ccmEFGH* genes represented *Nitrospira lenta* species and these genes were present in *Nitrospira lenta* reference genome, which was consistent with the taxonomy associated with the SSU rRNA gene in the *Nitrospira*-like MAG (C51).

In addition to the highly abundant nitrite oxidoreductase genes (*nxrAB*) (NODE_68), this dominant *Nitrospira*-like MAG (C51) had the potential for other nitrogen transforming reaction as it contained genes involved in assimilatory nitrate and nitrite reduction (*nasA* and *nirA*, respectively) as well as the nitric oxide forming nitrite reductase (*nirK*) (Fig. 6). This suggests that this MAG has the potential for complete assimilatory nitrate reduction to ammonia (i.e., *nasA* and *nirA*).

Correlations with other dominant MAGs revealed that within RES2, *Nitrospira*-like MAG (C51) had a strong negative correlation with *Nitrosomonas*-like MAG (C107) (Spearman correlation: −0.98, p < 0.001). Interestingly, *Nitrospira*-like MAG (C51) showed a strong negative correlation to another *Nitrospira*-like MAG (C56) in RES1 (Spearman correlation: - 0.79, p < 0.05), although this other *Nitrospira*-like MAG (C56) was present at very low coverage across all samples. Additional correlation analyses revealed strong negative correlations between *Nitrospira*-like MAG (C51) and *Sphingomonas*-like MAG (C70) and *Rhizobiales*-like MAG (C103.2) (C70: Spearman correlation: −0.73, p < 0.05 and C103.2: Spearman correlation: −0.64, p < 0.05) (Fig. S8).

#### 3.5.3 *Rhizobiales*-like MAG (C103.2)

The *Rhizobiales*-like MAG (C103.2) showed high coverage across both RES1 (25 ± 12 rpkm) and RES2 (20 ± 10 rpkm) constituting the second most abundant MAG across all samples (Fig. 7). The genetic potential of this MAG in terms of the nitrogen cycle included assimilatory nitrate and nitrite reduction (*nasA* and *nirAB*, respectively) also indicating the potential of this MAG for complete assimilatory nitrate reduction to ammonia (i.e., *nasA* and *nirA*) (Fig. 6). The poor taxonomic resolution of this MAG could not be improved based on the contig containing the SSU rRNA (NODE_1241) as it was also only identified as *Rhizobiales*. However, BLAST results of the *nasA* and *nirAB* genes revealed a close relation to *Proteobacteria* bacterium ST_bin 15 reported by Wang *et al*. (2017). Furthermore, assessment of additional correlations between *Rhizobiales*-like MAG (C103.2) and other dominant MAGs showed a strong positive correlation with *Sphingomonas*-like MAG (C70) (Spearman correlation: 0.76, p < 0.05) (Fig. S8).

#### 3.5.4 *Sphingomonas*-like MAG (C70)

Lastly, *Sphingomonas*-like MAG (C70) showed consistent coverage across both reservoirs (i.e. RES1: 11 ± 18 rpkm and RES2: 12 ± 18 rpkm) constituting the fourth most abundant MAG within the community (Fig. 7). However, the genetic potential for the involvement of this MAG in nitrogen transformations was limited as it was observed to only contain the nitric oxide reductase *norB* gene (Fig. 6). This corresponds to the increased number of *norB* genes identified (through BLAST analysis) as members of the family *Sphingomonaceae*. Further taxonomic classification of both the SSU rRNA contig and *norB* gene identified in this MAG, confirmed its taxonomy as *Sphingomonas*. The dominance of this MAG within the community suggests that it may be involved in other important metabolic capabilities within the microbial community that extend beyond the nitrogen cycle, such as biofilm formation.

## 4. Discussion

This study represents an in-depth investigation into the microbial nitrogen metabolism in chloraminated drinking water reservoirs. In doing so, we provide insight into the genetic potential of a chloraminated drinking water microbial community to transform nitrogen species and identify the dominant members responsible for specific pathways within the nitrogen cycle. Previous studies have generally concentrated on the relationship between chloramination and nitrification (Regan *et al*., 2003; Zhang *et al*., 2009; Sawade *et al*., 2016; Moradi *et al*., 2017). However, this study comprehensively explores nitrogen metabolism, where changes in the microbial community composition as well as the genetic potential in terms of nitrogen metabolism is revealed in two linked chloraminated reservoirs.

### 4.1 Varying stages of nitrification between the two reservoirs

The chemical monitoring data indicated differing patterns in the concentrations of nitrogen species and spatial changes in disinfectant residuals in the two reservoirs, suggesting different nitrification regimes in the two reservoirs. Varying stages of nitrification in different sections of a chloraminated DWDS was also previously observed by Shaw *et al*. (2015). Nitrification was observed, specifically in RES2, as decreases in ammonium concentrations were associated with concomitant increases in nitrite and nitrate concentrations. An increase in the concentrations of both nitrite and nitrate were observed following RES1, although their concentrations were low and did not consistently correlate to changes in ammonium concentrations. While nitrification occurred in both reservoirs, it did not occur to a significant extent in RES1 for all sampled time points. This may be a consequence of increased monochloramine concentrations, as months with increased monochloramine concentrations showed reduced nitrification rates. In addition, ammonium concentrations were consistently higher in RES1 compared to RES2, as expected as these samples were closer to the site of chloramination. Conversely, following RES2, samples typically had elevated nitrate concentrations with reduced nitrite and ammonium concentrations, suggesting that with increasing distance from the site of chloramination, samples nearing the end of the DWDS undergo complete nitrification. Here it is more likely that the depletion of nitrite may be the result of tight coupling of canonical ammonia and nitrite oxidation (Shaw *et al*., 2015; Daims *et al*., 2016).

### 4.2 Changes in bacterial community composition between the two reservoirs

The majority of annotated proteins were bacterial and the taxonomic profiles based on SSU rRNA genes were in agreement with previous descriptions of the microbial community within chloraminated DWDS, in which *Proteobacteria* and *Nitrospira* (to a lesser extent) were the dominant phyla (Bautista-de los Santos *et al*., 2016; Gülay *et al*., 2016: Potgieter *et al*., 2018). The change in dominance between *Alphaproteobacteria* and *Betaproteobacteriales* in RES1 and RES2 was also observed by Potgieter *et al*., 2018, where *Betaproteobacteriales* were more abundant closer to the site of chloramination and an increase in *Alphaproteobacteria* correlated to decreased disinfectant residual concentrations in RES2. The variation in dominance between *Alphaproteobacteria* and *Betaproteobacteria* (now classified as the order *Betaproteobacteriales* in the phylum *Gammaproteobacteria*) has been well documented, where their dominance varies depending on multiple factors including disinfectant residual concentrations (Gomez-Alvarez *et al*., 2012; Wang *et al*., 2014) and seasonal trends (McCoy and VanBriesen, 2014; Pinto *et al*., 2014; Prest *et al*., 2016; Zlatanovic *et al*., 2017). Therefore, in this study, the increased retention time in the reservoirs may have an effect on the microbial community composition and function in terms of nitrogen transformation.

However, due to the lack of quantitative and viability assays it is unclear what proportion of the community data is from viable or active cells. Although there is a lack of absolute abundance in this study, it does not detract from the observed genetic potential of the microbial community to transform nitrogen. Furthermore, the coverage of the highly abundant MAGs (specifically, *Nitrosomonas, Sphingomonas* and *Nitrospira*) corresponds to that found by Sakcham *et al*. (2019) where after removal of eDNA, *Nitrosomonas* was present at greater coverage than *Nitrospira*. Furthermore, at a location with higher nitrite concentrations and after eDNA removal, *Sphingomonas* showed and increase in relative abundance, which correlated to higher abundances of *Nitrosomonas* at this location. This positive correlation between *Sphingomonas* and *Nitrosomonas* was also observed in this study. Here, *Sphingomonas* may act as a potential indicator for the onset of nitrification in chloraminated systems (Sakcham *et al*., 2019).

### 4.3 Nitrification driven by co-occurring *Nitrosomonas* and *Nitrospira* species

The engineered oligotrophic environment of drinking water systems typically have limited substrate for nitrifiers. However, in chloraminated drinking water systems, nitrogen (in the form of ammonia) is available either as excess ammonia or through disinfectant decay. This addition of ammonia consequently promotes the growth of nitrifying bacteria and drives nitrification (Regan, 2003; Wang *et al*., 2017). The phylogenetic distribution of most commonly occurring ammonia-oxidising bacteria and archaea has been well described with AOB falling within *Gammaproteobacteria* (order: *Betaproteobacteriales*), the AOA falling specifically within the *Thaumarchaea* (Pester *et al*., 2011). However, in this study *Thaumarchaeota* was not detected. AOA are thought to play a more significant role in nitrification in environments that are low in dissolved oxygen or low ammonia concentrations (Schleper *et al*., 2010; Pester *et al*., 2011). In contrast, *Betaproteobacteriales* AOB, specifically *Nitrosomonas* were present in high abundances, indicating that *Nitrosomonas* is a dominant member of the microbial community and plays a significant role in the fate of nitrogen in this chloraminated system. The finding of AOB dominance over AOA is consistent with other chloraminated DWDS studies (de Vet *et al*., 2011; Wang *et al*., 2014).

Nitrification in chloraminated drinking water systems has been well characterised with *Nitrosomonas* and *Nitrospira* species often identified as the major ammonia- and nitrite-oxidisers, respectively (Regan, 2003; Hoefel *et al*., 2005; Zhang *et al*., 2009). In this study, the dominance of *Nitrosomonas* and *Nitrospira* species was confirmed as these species showed: (i) high abundance of their respective SSU rRNAs, (ii) high coverage of nitrifying genes associated with *Nitrosomonas* (*amoABC* and *hao* genes) and *Nitrospira* (*nxrAB* genes), (iii) high coverage of their respective *Nitrosomonas* and *Nitrospira* reference genomes and lastly (iv) high coverage of constructed *Nitrosomonas*-like and *Nitrospira*-like MAGs, across both reservoirs. The dominance of *Nitrosomonas* species in this study corresponded to the high abundance of *Nitrosomonas* observed by Potgieter *et al*. (2018), where they observed *Nitrosomonas*-like OTUs to be the most abundant OTUs, constituting 18.3% of the total sequences. Furthermore, *Nitrospira* was identified as the second most abundant phyla in this study, which correlated to the observed increase of *Nitrospira* in the reservoir samples observed by Potgieter *et al*. (2018).

Here, the chemical monitoring data alone may be limited in conclusively confirming the occurrence of nitrification however, the increased concentrations of nitrate together with the high abundance of *Nitrosomonas* and *Nitrospira* strongly suggests that nitrification occurs within both chloraminated reservoirs. Furthermore, Duff *et al*. (2017) reported that *amoA* gene abundances showed significant correlations to the Potential Nitrification Rate (PNR) from AOB in marine inertial bays. Here the high abundance of genes involved in ammonia-oxidation correlates to the observed increase in nitrate concentrations.

The phylogeny of the *amoA* gene confirmed that the majority of AOB were identified as strict ammonia oxidisers (specifically *Nitrosomonas*) with the presence of one potential comammox *amoA* distinct from the canonical AOBs. Some AOB representatives, such as *Nitrosomonas oligotropha*, may have a higher affinity for ammonia (Regan, 2003). Given the low concentrations of ammonia in these systems, the high affinity of *Nitrosomonas oligotropha* may allow it to outcompete other *Nitrosomonas* species (Regan, 2003). This is likely to be the case in RES2 where ammonium concentrations are lower than in RES1. Here, where the *Nitrosomonas*-like MAG (C107) decreases in coverage, the other *Nitrosomonas*- like MAG (C58) increases, indicating that C58 may have a higher affinity for ammonia when ammonia concentrations are low and is therefore able to outcompete other *Nitrosomonas*-like population.

Similarly, the phylogeny of *nxrA* genes showed that the majority of *nxrA* genes group within *Nitrospira* lineage II, which is the most widespread and diverse of the lineages, including both canonical NOBs as well as all currently known comammox bacteria (Koch *et al*., 2015; Daims *et al*., 2016; Daims and Wagner, 2018). The majority of *nxrA* genes from this study grouped closely with *Ca*. Nitrospira sp. ST-bin5 described by Wang *et al*. (2017) and the grouping of *Ca*. Nitrospira sp. ST-bin5 with *Nitrospira moscoviensis* was observed in both this study and the study by Wang *et al*. (2017). Furthermore, in their study they describe that *Ca*. Nitrospira sp. ST-bin5 does not contain the genes responsible for ammonia oxidation (*amo* and *hao*) and is therefore not a true comammox bacteria. However, it is not possible to identify the presence of comammox bacteria based on *nxrA* phylogeny.

It is now known that comammox and canonical AOB both use ammonia as substrate and therefore may co-exist in niches with low ammonia concentrations such as drinking water (Wang *et al*., 2017). Therefore, investigations into the presence of comammox revealed potential comammox based on *amoA* phylogeny and the identification of cytochrome c biogenesis genes (*ccmEFGH*) in both *Nitrospira*-like MAGs (Palomo *et al*., 2018). However, *ccmEFGH* genes were found to belong to *Nitrospira lenta* species, which was consistent with the taxonomy of the SSU rRNA within the *Nitrospira*-like MAG (C51). In addition, cyanate hydratase or cyanase (*cynS*) and assimilatory nitrite reductase (*nirA*) genes were observed in both *Nitrospira*-like MAGs. These genes are typically absent in comammox and only detected in canonical NOB *Nitrospira*, which further confirms that the *Nitrospira*-like MAG (C51) is most likely related to *Nitrospira lenta*.

Furthermore, the dominance of one *Nitrospira*-like MAG (C51) was observed over another *Nitrospira*-like MAG (C56) indicating its ability to outcompete other *Nitrospira* members similar to what has been demonstrated in activated sludge systems (Maixner *et al*., 2006; Ushiki *et al*., 2017). Competition and separation in ecological niches between *Nitrospira* may be caused by physiological properties such as the affinity for nitrite and other substrates, formate utilisation and the relationship with AOB (Ushiki *et al*., 2017). Typically, nitrification occurs in a modular fashion performed by a complex network of specialised microorganisms. This modularity results in the cooperative and competitive interactions (Stein and Klotz, 2011; Kuypers *et al*., 2018). Nitrification generally occurs through a cooperative interaction between ammonia and nitrite oxidisers. Competitive interactions may exist within the ammonia-oxidising and nitrite-oxidising groups, which potentially compete for substrates ammonia and nitrite, respectively (Kuypers *et al*., 2018). Here, the dominant *Nitrospira*-like MAG (C51) showed a positive relationship with the *Nitrosomonas*-like MAG (C107), decreasing in coverage in RES2 where disinfectant residuals and consequently ammonium concentrations were lower. This relationship may involve the tight coupling of canonical ammonia and nitrite oxidation, thereby resulting in the low concentrations of nitrite observed in both reservoirs (Shaw *et al*., 2015; Daims *et al*., 2016).

### 4.4 Genetic potential of the microbial community for nitrogen metabolism

Despite the limitations in available drinking water metagenomes, the results in this study provide insight into the diversity of microorganisms involved in nitrogen transformation in chloraminated drinking water. As a consequence of nitrification, concentrations of nitrate increase, thereby providing an alternative form of biologically available nitrogen. The increased availability of nitrate potentially promotes the growth of a highly diverse assemblage of microorganisms. There is an astounding diversity of microorganisms that transform nitrogen, where each microorganism has specific physiological requirements for optimal growth. Nitrogen transformations in the environment are typically carried out by microbial communities that recycle nitrogen more efficiently than a single microorganism. These microbial communities retain their nitrogen transforming capabilities even when the community composition is altered with changes in the environment. These microbial nitrogen transforming reactions form complex networks in both the natural and engineered environments (Kuypers *et al*., 2018).

Where nitrification is driven by a select few microorganisms (i.e., *Nitrosomonas* and *Nitrospira*), the reduction of nitrate, nitrite and nitric oxide was likely performed by a diverse assembly of bacteria (predominately *Alphaproteobacteria*, *Gammaproteobacteria*, order *Betaproteobacteriales* and *Nitrospira*), each with their own discrete physiological requirements for optimal growth (Kuypers *et al*., 2018). The dominance of *amoABC* and *hao* (ammonia oxidation), *nxrAB* (nitrite oxidation), *nasA* (assimilatory nitrate reduction), *nirBD* (nitrite reductase), *nirK* (nitrite reductase, NO-forming) and *norBCDQ* (nitric oxide reduction) genes suggests that ammonia is oxidised and the resulting in the formation of nitrite and nitrate. The resulting oxidised compounds are most likely either (i) assimilatoraly reduced to nitric oxide and ultimately nitrous oxide potentially triggering biofilm formation or (ii) when ammonia concentrations are low, nitrite and nitrate are fixed and converted to ammonia for assimilation (Fig. 8).

**Fig. 8:**
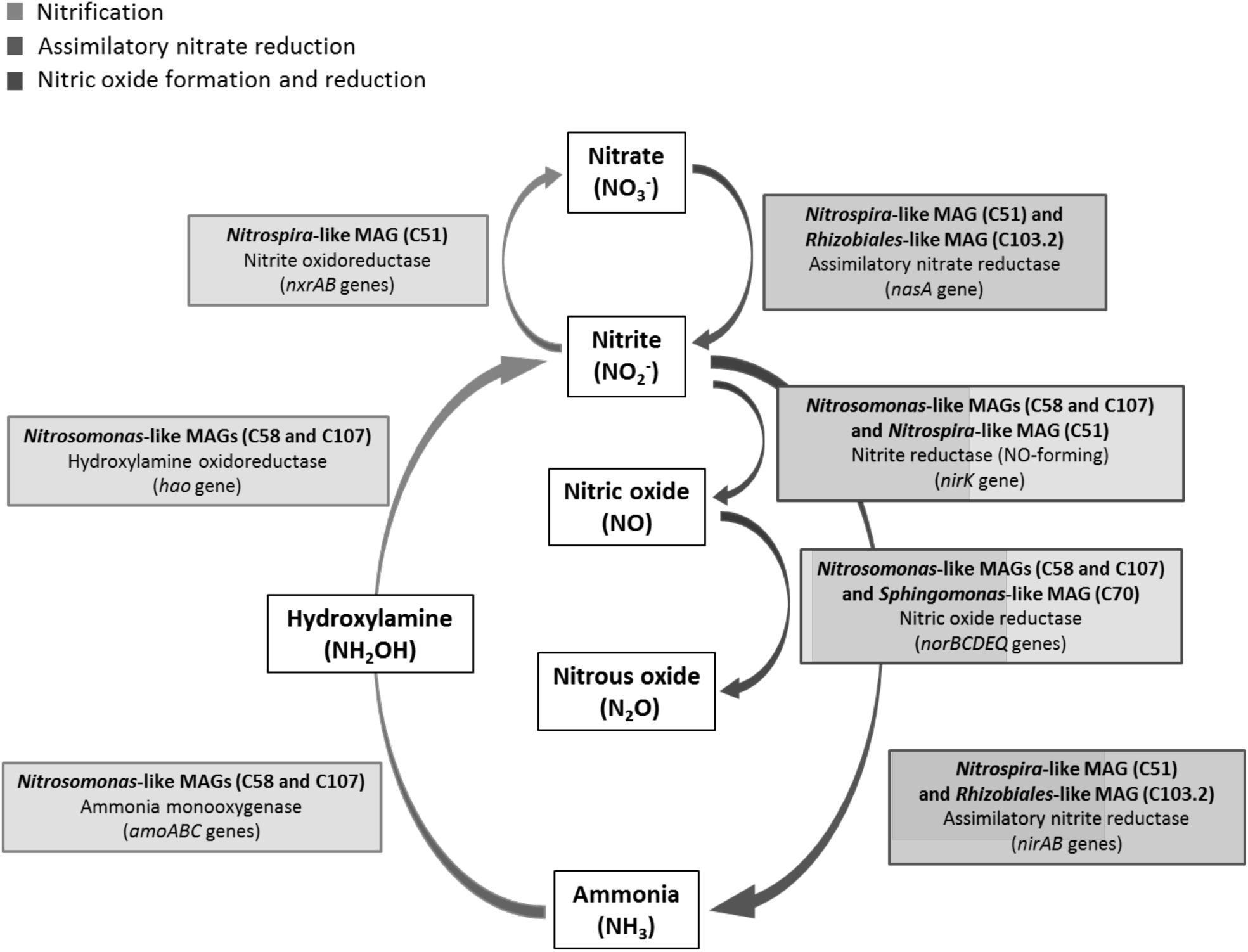
A schematic overview of the dominant genes and MAGs involved in the major reactions of the nitrogen cycle both chloraminated reservoirs. Reactions involved in nitrification are indicated in green, reactions involved in assimilatory nitrate reduction are indicated in blue and the reactions involving the formation and reduction of nitric oxide are indicated in red.

Bacterial nitrate reduction has been shown to be a multifaceted process, performed by three distinct classes of nitrate reducing systems differentiated by their cellular location, regulation, structure, chemical properties and gene organisation (Moreno-Vivian *et al*., 1999). In this study, all three systems (i.e., Nas, Nar and Nap) were observed within the diverse group of recovered draft genomes. The diversity observed within nitrate reductases in this study correlated with that observed in other studies where nitrate reductases were found to phylogenetically widespread (Richardson *et al*., 2001; Philippot, 2005; Smith *et al*., 2007). In many cases in this study, a single draft genome contained genes for both assimilatory and dissimilatory nitrate reduction, indicating that these pathways may be interconnected where the enzymes may play different roles under different metabolic conditions (Moreno-Vivian *et al*., 1999). The interconnection of assimilatory, respiratory and dissimilatory nitrate reduction may facilitate rapid adaptation to changes in nitrogen and/or oxygen conditions thereby increasing the metabolic potential for survival in the chloraminated drinking water environment (Moreno-Vivian *et al*., 1999).

In this study, the potential for assimilatory nitrate reduction was the dominant nitrate reducing pathway as *nasA* genes were highly abundant and were the only nitrate reductase genes observed in the dominant MAGs, i.e. the *Nitrospira*-like MAG (C51) and the *Rhizobiales*-like MAG (C103.2). Assimilatory nitrate reductases are typically cytoplasmic and enables the utilisation of environmental nitrate as a nitrogen source. This enzyme is generally induced by nitrate but inhibited by ammonium, however it is not affected by oxygen (Moreno-Vivian *et al*., 1999). This converse relationship was observed between the abundance of *nasA* genes and ammonia concentrations. The high abundance of genes associated with assimilatory nitrate reduction in this study correlated with reduced concentrations of ammonium and increased concentrations of nitrate, which is a result of nitrification. In the oligotrophic environment of DWDS, where nutrients are limited, utilization of nitrate as nitrogen source for biomass synthesis may be an important mechanism for survival (Rivett *et al*., 2008).

The metabolic fate of nitrite formed through the reduction of nitrate may involve subsequent reduction to ammonia (through assimilatory nitrite reduction) or to nitric oxide (Moreno-Vivian *et al*., 1999; Smith *et al*., 2007; Daims *et al*., 2016). Both *nirA* and *nirB* genes encoding nitrite reductases were observed to have high coverage across both reservoirs. The positive correlation with *nasA* and *nirA* confirms the potential for complete assimilatory nitrate reduction to ammonia, specifically in RES2. However in RES1, a converse relationship was observed between *nasA* and *nirA* genes, suggesting that *nirB* may potentially may play a larger role in the reduction of nitrite here as *nirB* genes showed similar trends to *nasA* genes in RES1, although, at a lower coverage than *nirA*. The NirB nitrite reductase uses NADH as an electron donor to reduce nitrite in the cytoplasm and typically, high nitrite concentrations are needed for *nirB*, which is consistent with the proposed role of the NirB enzyme in detoxification of nitrite. This may be the case following RES1, where nitrite concentrations were generally higher as compared to RES2.

Genes encoding dissimilatory nitrate reduction and denitrification were identified in this study, however their coverage across both reservoirs was generally very low. Both dissimilatory nitrate reduction to ammonia (DNRA) and denitrification compete for the nitrate and nitrite as an electron acceptor (Smith *et al*., 2016). It has been shown that DNRA may be favoured when nitrate concentrations are low and organic electron donor availability is high, whereas denitrification outcompetes DNRA when nitrate concentrations are high and carbon supplies are limiting (Rivett *et al*., 2008; Smith *et al*., 2016). However, denitrification typically occurs in environments where available oxygen is limited and nitrate is used in respiration. Therefore, denitrification may not play a role in an aerobic drinking water environment.

Alternatively, nitrite may be reduced to nitric oxide (NO) via *nirK* (nitrite reductase, NO-forming), which showed high coverage across both reservoirs and was observed in *Nitrosomonas*-like and *Nitrospira*-like MAGs. At low, non-toxic concentrations, NO can potentially elicit other cellular responses other than denitrification (Arora *et al*., 2015). Although it is difficult to separate NO signalling responses from detoxification and denitrification, in *Nitrosomonas europaea* it was observed that low concentrations of NO caused biofilm dispersal, whereas high concentrations of NO caused increased biofilm formation as a defence mechanism (Arora *et al*., 2015). In this study, generally a converse relationship was observed between assimilatory nitrite reduction (*nirAB*) and NO forming nitrite reduction (*nirK*) suggesting that potentially when ammonia concentrations are increased, the need for nitrite to be assimilated to ammonia is reduced and nitrite may be preferentially reduced to nitric oxide. Furthermore, *nor* genes were observed to have high coverage across reservoir samples and were present in both *Nitrosomonas*-like MAGs and the *Sphingomonas*-like MAG. *Nor* genes are responsible for regulating the concentration of NO by the reduction of NO to nitrous oxide (N_2_O). As denitrification may not be an important process in this system and the observed high coverage of *nirK* and *nor* genes, suggests that production of NO may act as an important molecule in regulation biofilm formation. This is consistent with *Sphingomonas* spp. being linked to the initiation of biofilm formation and resistance to monochloramine (Chiao *et al*., 2014).

#### 4.4.1 Other nitrogen transforming processes

The metabolic versatility of nitrogen transforming processes included nitrogen fixation. Although *nif* genes were observed at very low coverages, a full cascade of *nif* genes were identified in a *Sideroxydans*-like MAG, which is well described as an iron-oxidising bacteria. *Sideroxydans*, has previously been described as commonly occurring OTU in disinfectant free drinking water (Bautista-de los Santos *et al*., 2016) however, the presence of these iron-oxidisers in DWDS are typically, linked to microbial mediated corrosion of the steel and iron pipes (Emerson and De Vet, 2015). Further investigations of the interactions between iron-oxidation and nitrogen fixation may be needed for the optimisation of the design and operations of chloraminated DWDS.

## 5. Conclusion

The transformation of nitrogen has been well characterised in many environments including the open oceans and ocean sediments, wastewater and agricultural soils. However, to our knowledge this information is limited when considering chloraminated DWDSs, where nitrification is a major concern. In this study, *Nitrosomonas* and *Nitrospira* were identified as the main drivers in nitrification and together with the water chemistry data (i.e., changes in ammonia, nitrite and nitrate concentrations), this study improves our understanding nitrogen cycling potential in chloraminated drinking water. Furthermore, the genes and bacteria responsible for the metabolic fate of nitrate and other nitrogen transforming reactions were identified and were shown to be highly diverse. Here, the addition of ammonia through chloramination promotes/supports the growth of nitrifying bacteria and in this DWDS, the nitrate formed through nitrification may ultimately be reduced via assimilatory processes either to ammonia when ammonia concentrations are low or to nitric oxide for potential regulation of biofilm formation. This study therefore provides insight into the genetic network behind microbially mediated nitrogen metabolism in chloraminated drinking water.

## Acknowledgements

This research was supported and funded by Rand Water, Gauteng, South Africa through the Rand water Chair in Water Microbiology at the University of Pretoria. Sarah Potgieter would also like to acknowledge the National Research Foundation (NRF) for additional funding. AJP was supported by National Science Foundation awards (CBET-1703089 and CBET-1749530). Furthermore, the authors would like to acknowledge the Agricultural Research council – Biotechnology platform (ARC-BTP) for their services in Illumina HiSeq shotgun sequencing as well as Holger Daims for the sharing of *nxrA* gene protein sequences from their Kitzinger *et al*. (2018) study.

## Supplemental material

**Fig. S1:** Spatial changes in the concentrations of nitrogen species

**Fig. S2:** Temporal changes in nitrogen compounds

**Fig. S3:** The relative abundance of bacterial SSU rRNA genes

**Fig. S4:** Alpha diversity measures of samples from both reservoirs

**Fig. S5:** The number of genes identified for each dominant function

**Fig. S6:** Temporal trends in the coverage of the dominant nitrogen transforming genes

**Fig. S7:** Log transformed coverage of the 47 MAGs

**Fig. S8:** Temporal trends in the coverage of the dominant MAGs

**Fig. S9:** Phylogenetic tree of *Nitrosomonas* SSU rRNA contigs

**Fig. S10:** Phylogenetic tree of *Nitrospira* SSU rRNA contigs

**Fig. S11:** Heatmap showing the coverage of reference genomes

**Table S1A:** Chemical monitoring data for RES1

**Table S1B:** Chemical monitoring data for RES2

**Table S2:** Chloraminated reservoir sample description

**Table S3:** The final number of reads mapped to assembly

**Table S4:** Genes identified in nitrogen transformation pathways

**Table S5:** Reference genomes of nitrifying organisms

**Table S6:** Characteristics of Metagenome Assembled Genomes (MAGs)

**Table S7:** Percentage coverage of SSU rRNA contigs identified as bacteria phyla

**Table S8:** Percentage coverage of SSU rRNA contigs identified as Eukaryota phyla

