## Supplemental figures and tables for "Microbial Nitrogen Metabolism in Chloraminated Drinking Water Reservoirs"

**Supplemental material**

**Figures**

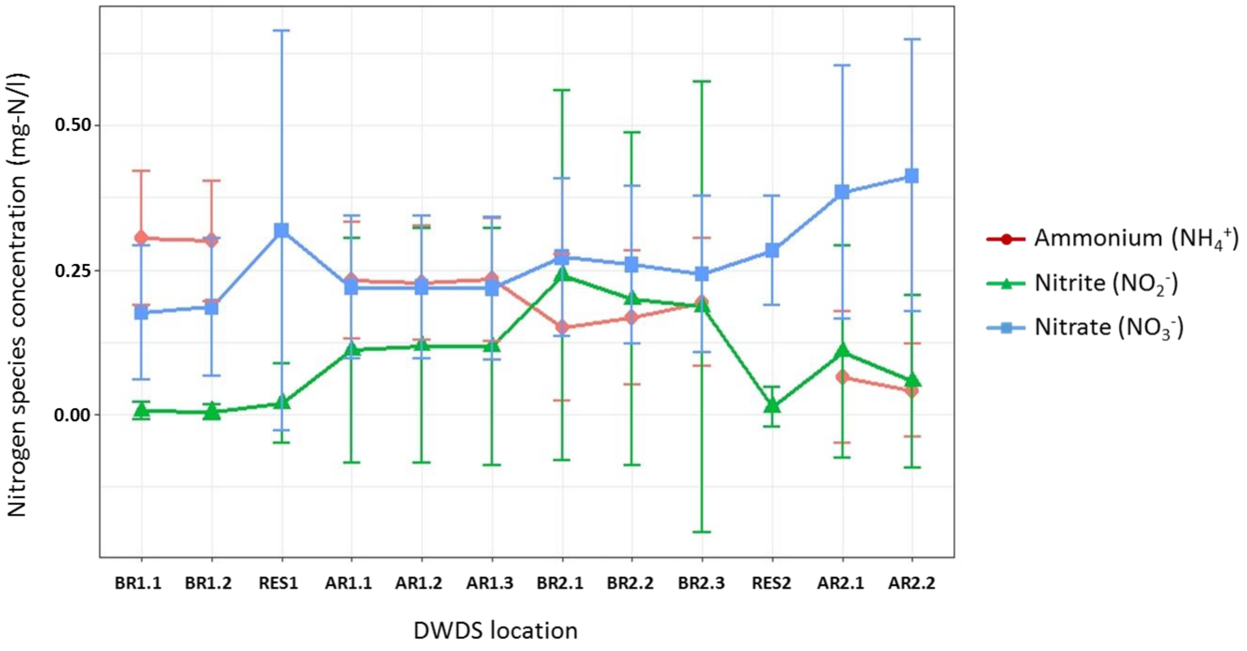


**Fig. S1**: Spatial changes in the concentrations of nitrogen species [i.e., ammonium ($NH_4^+$), nitrite ($NO_2^-$) and nitrate ($NO_3^-$)] for locations within the chloraminated section of the DWDS averaged over two years. Error bars capture variation in concentration of each nitrogen species over the period of two years. Location names on the x-axis translate to: before reservoir 1 (BR1.1 and BR1.2), reservoir 1 (RES1), after reservoir 1 (AR1.1 – AR1.3), before reservoir 2 (BR2.1 – BR2.3), reservoir 2 (RES2) and locations after reservoir 2 (AR2.1 and AR2.2). Ammonium concentration data is missing for sites RES1 and RES2.

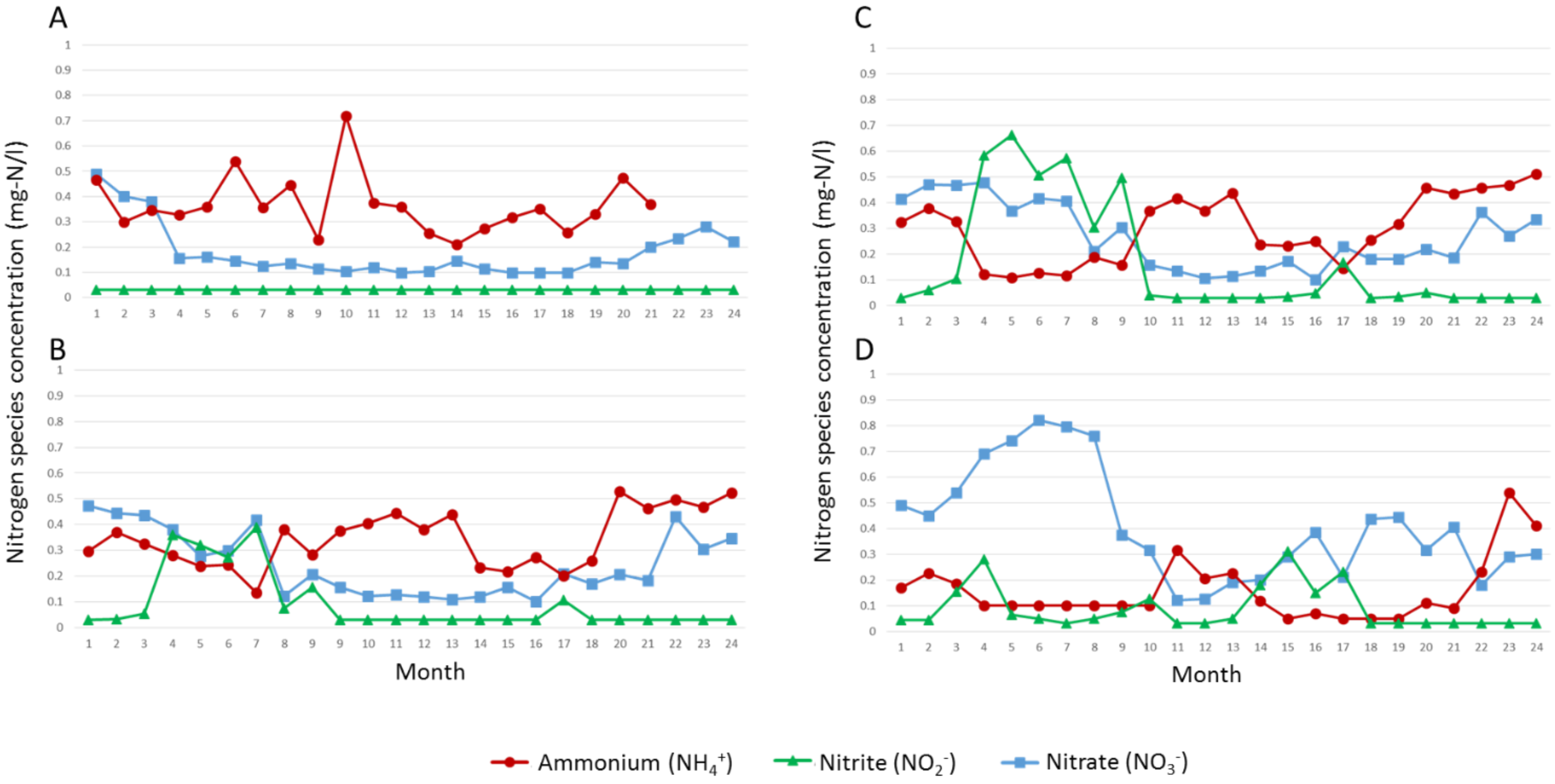


**Fig. S2**: Temporal changes in nitrogen compounds [i.e., ammonium (red circles), nitrite (green triangles) and nitrate (blue squares)] over 2 years. Plots (A) and (B) refer to before and after RES1, respectively and plots (C) and (D) refer to before and after RES2, respectively.

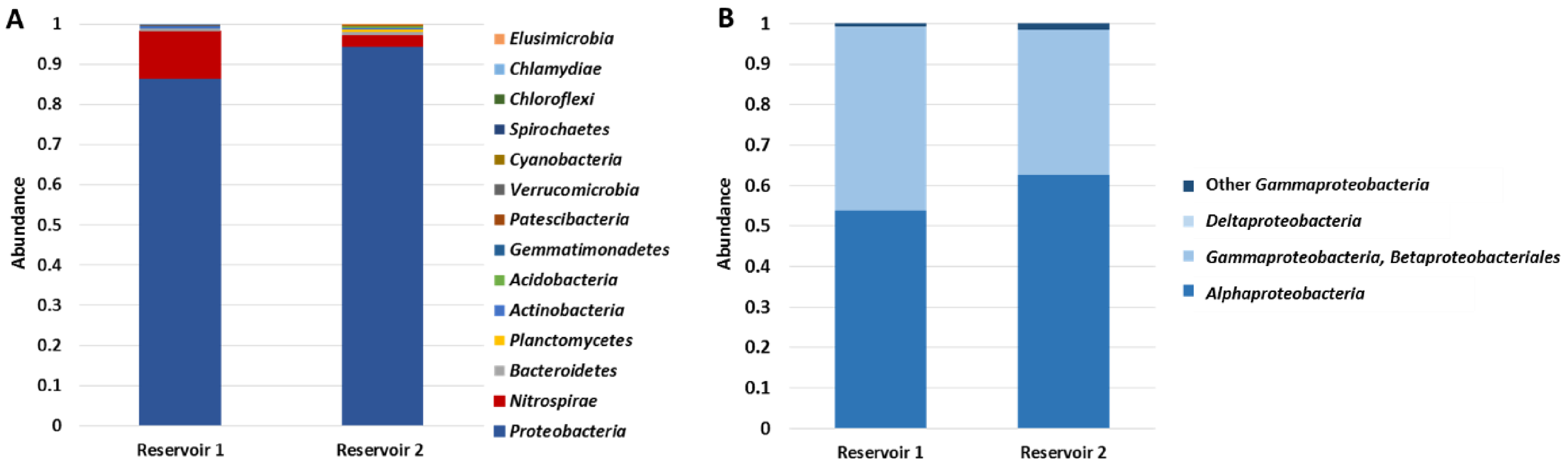


**Fig. S3**: The relative abundance of bacterial SSU rRNA genes i.e., bacterial phyla (A) and proteobacterial classes (B). The relative abundance of SSU rRNA genes within each taxonomic grouping was averaged across all samples for each reservoir (RES1 and RES2) (Coverage calculated for contigs > 250bp).

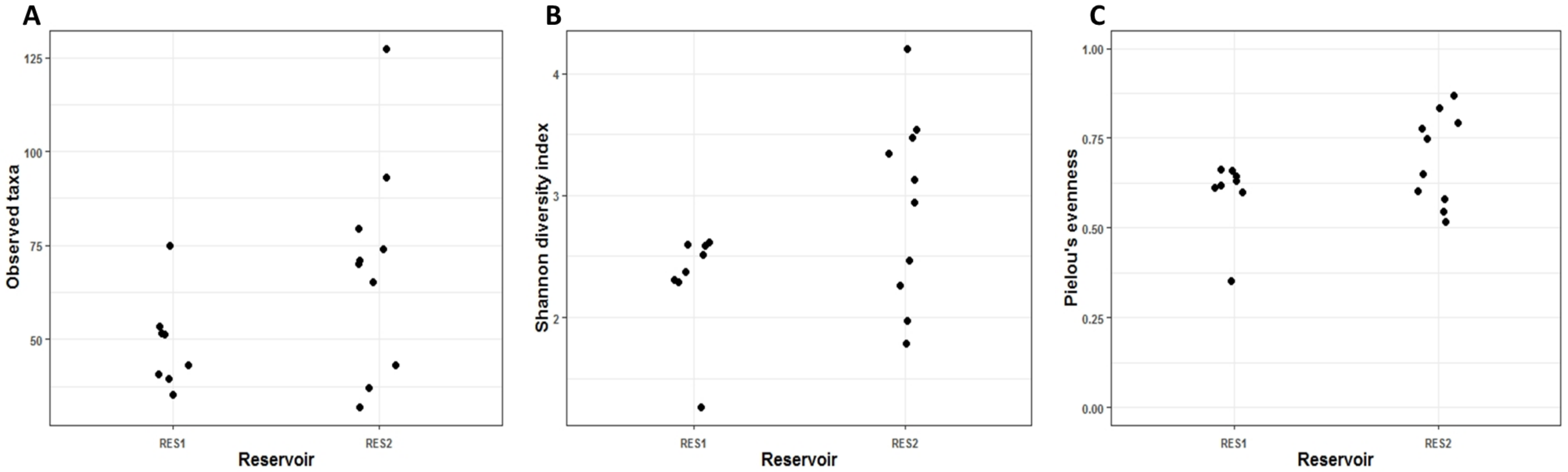


**Fig. S4**: Alpha diversity measures of samples from both reservoirs (RES1 and RES2) i.e. richness (A), Shannon diversity (B) and evenness (C).

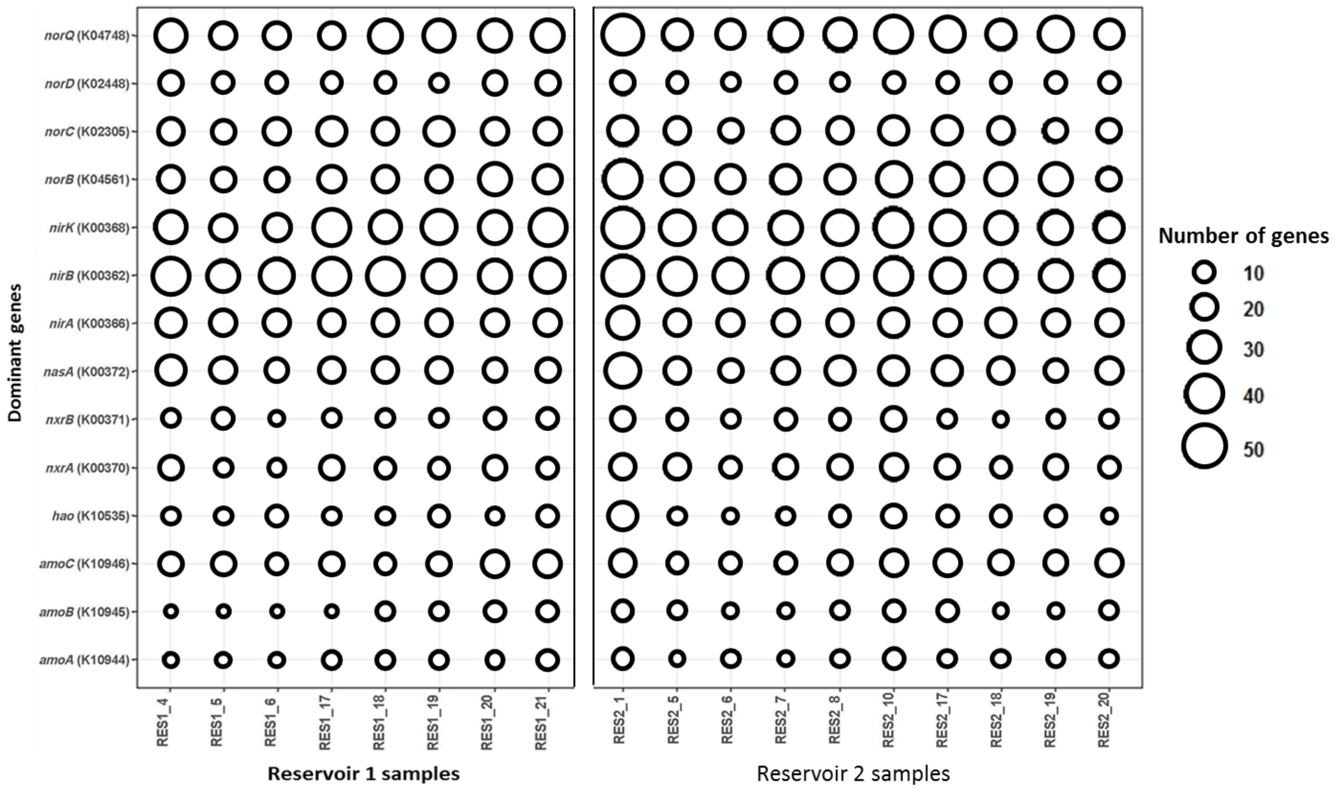


**Fig. S5**: The number of genes identified for each dominant function across all reservoir samples.

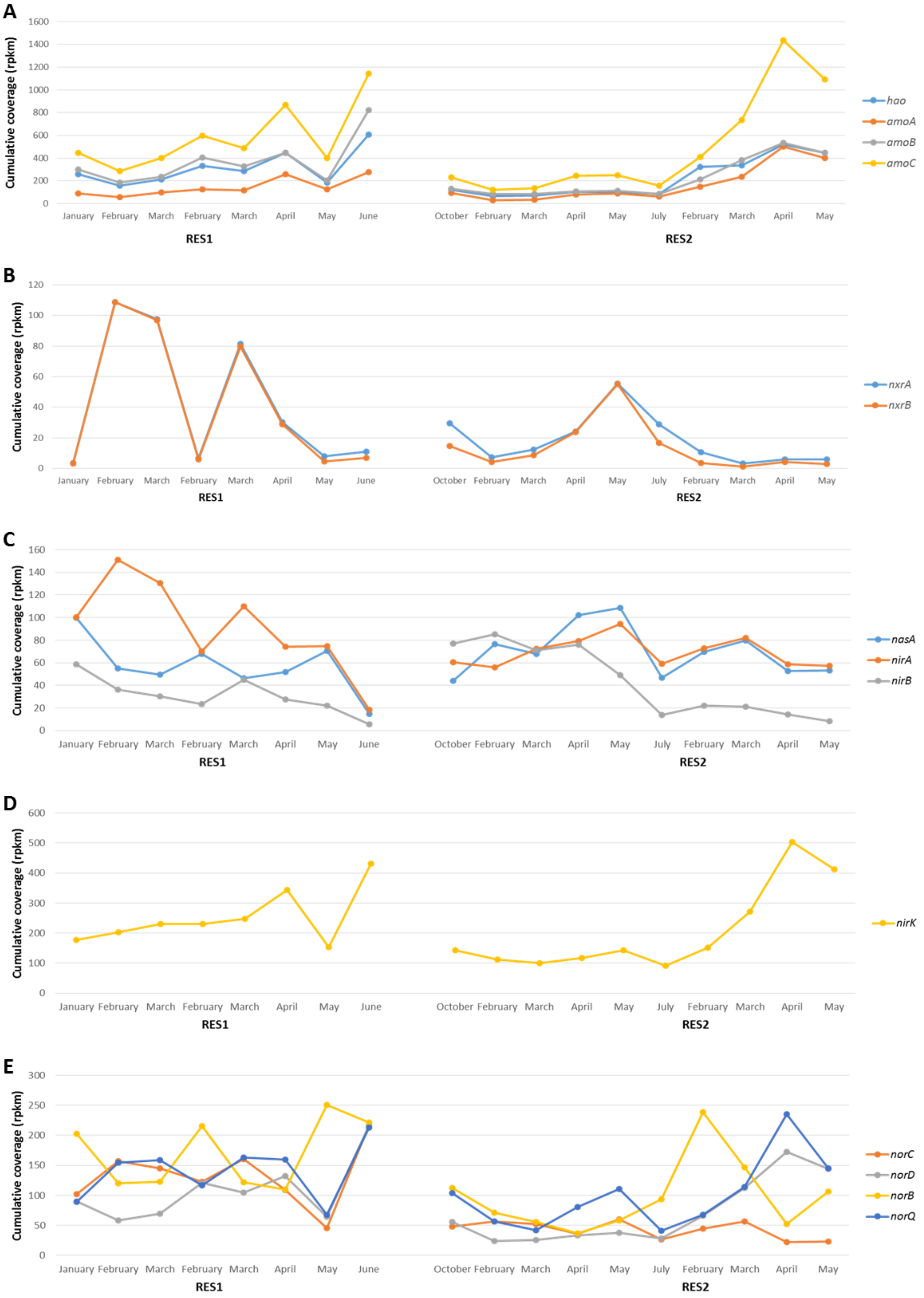


**Fig. S6**: Temporal trends in the cumulative coverage of the dominant nitrogen transforming genes identified in the dataset. (A) ammonia oxidation (*amoABC* and *hao* genes), (B) nitrite oxidation (*nxrAB* genes), (C) assimilatory nitriate and nitrite reduction (*nasA* and *nirAB* genes), (D) nitrite oxidation, nitric oxide-forming (*nirK* gene) and (E) nitric oxide reduction (*norBCDQ* genes).

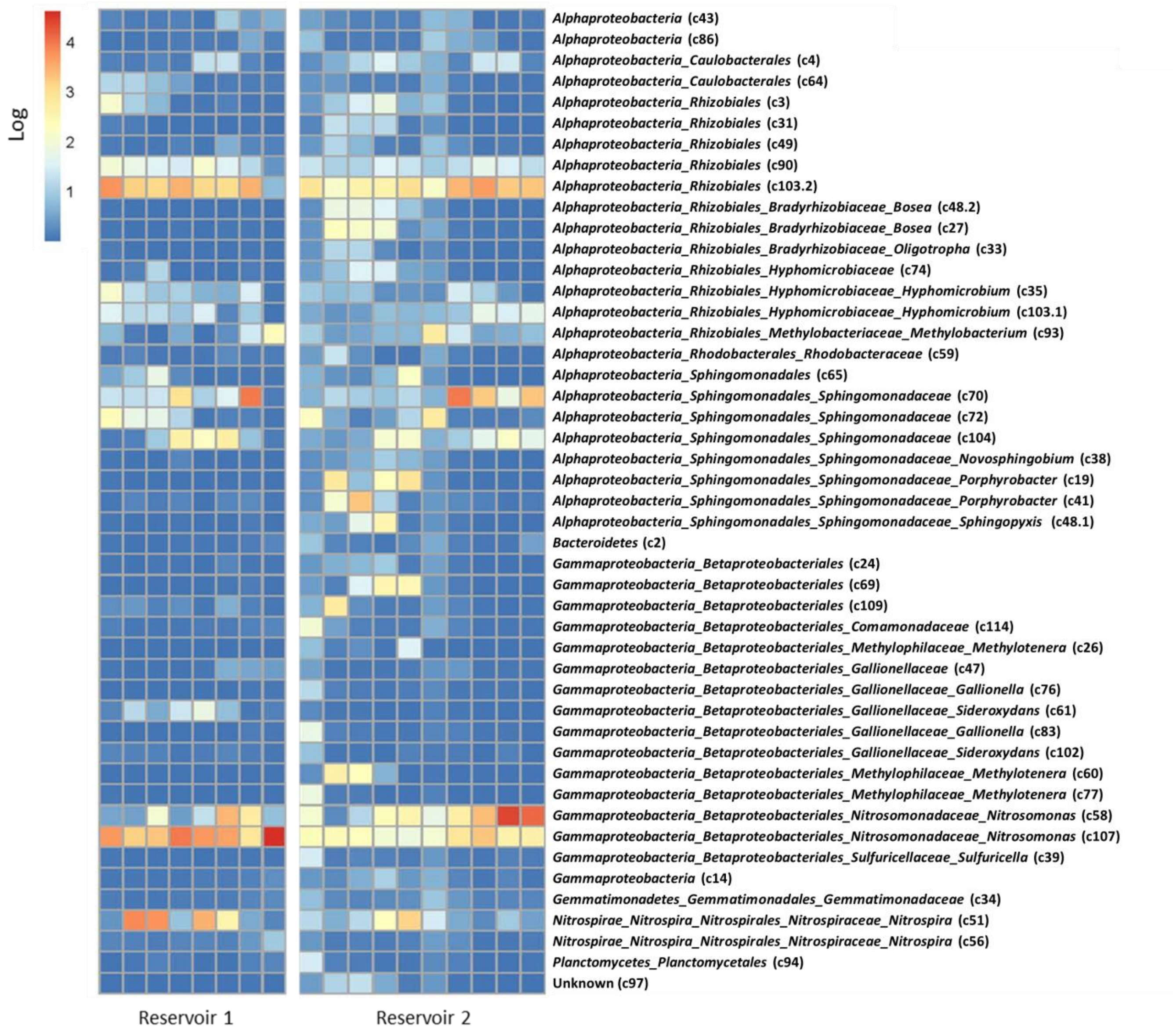


**Fig. S7**: Log transformed coverage of the 47 Metagenome Assembled Genomes (MAGs) across samples from the two reservoirs. The level of coverage is indicated by the key on the top left of the Fig. as the log of reads per kilo base million (rpkm).

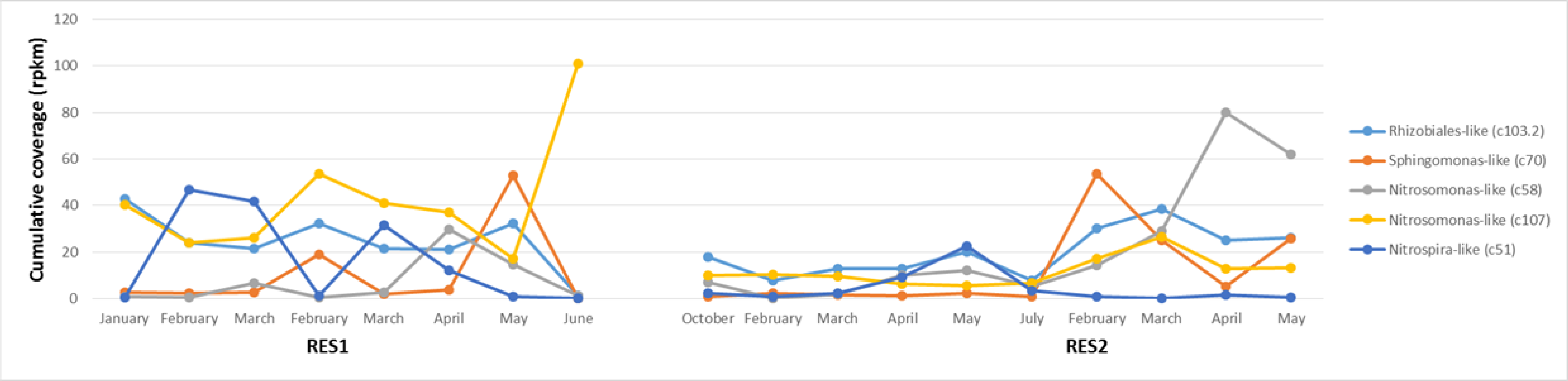


**Fig. S8**: Temporal trends in the cumulative coverage of the dominant Metagenome Assembled Genomes (MAGs) constructed in the dataset.

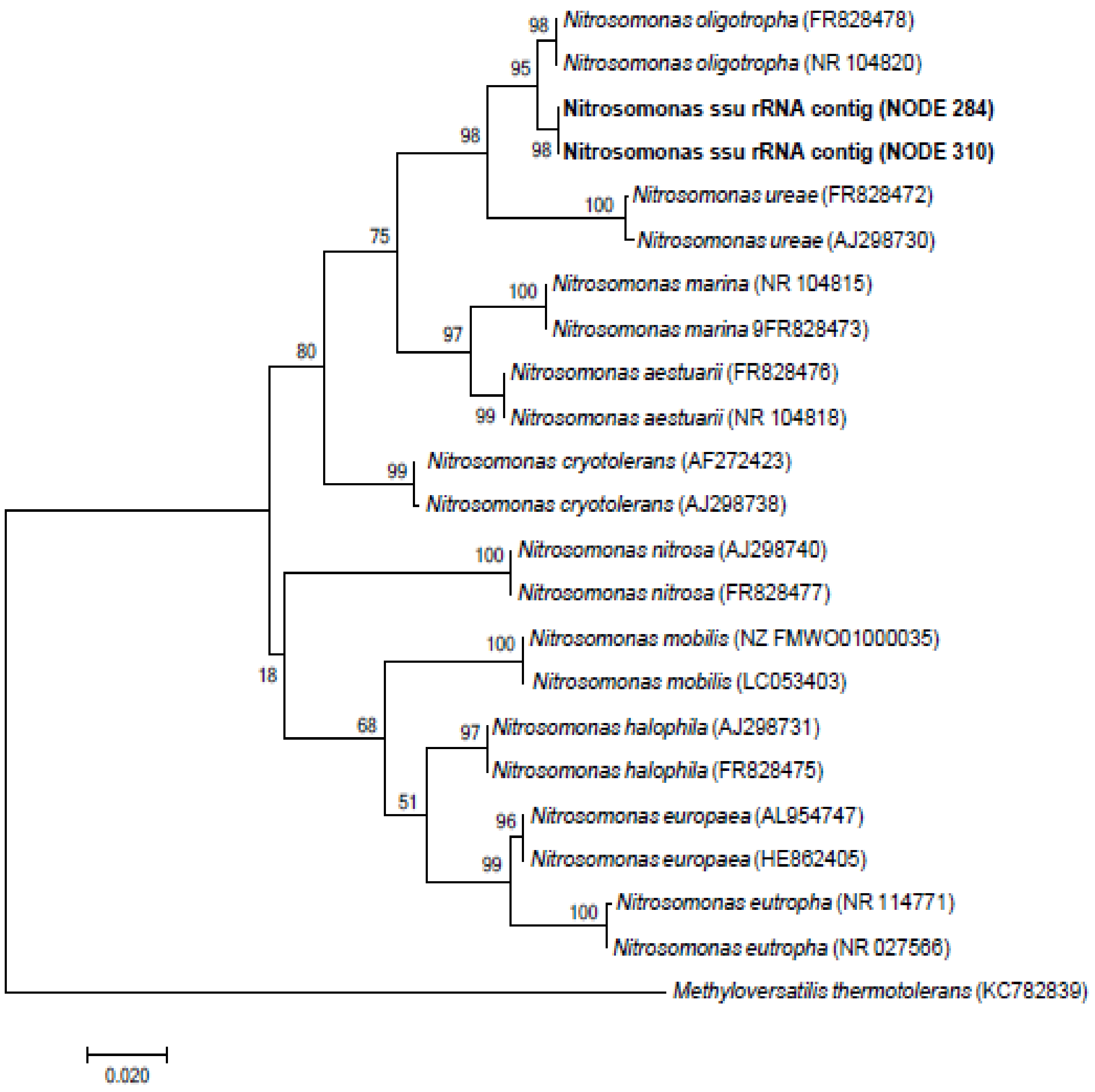


**Fig. S9**: Maximum likelihood phylogenetic tree showing the grouping of *Nitrosomonas* SSU rRNA contigs retrieved from metagenomic data with *Nitrosomonas* spp. 16S rRNA reference strains. With *Methyloversatilis thermotolerans* as the outgroup and bootstrap analysis of 1000 replicates (bootstrap values are indicated as percentages).

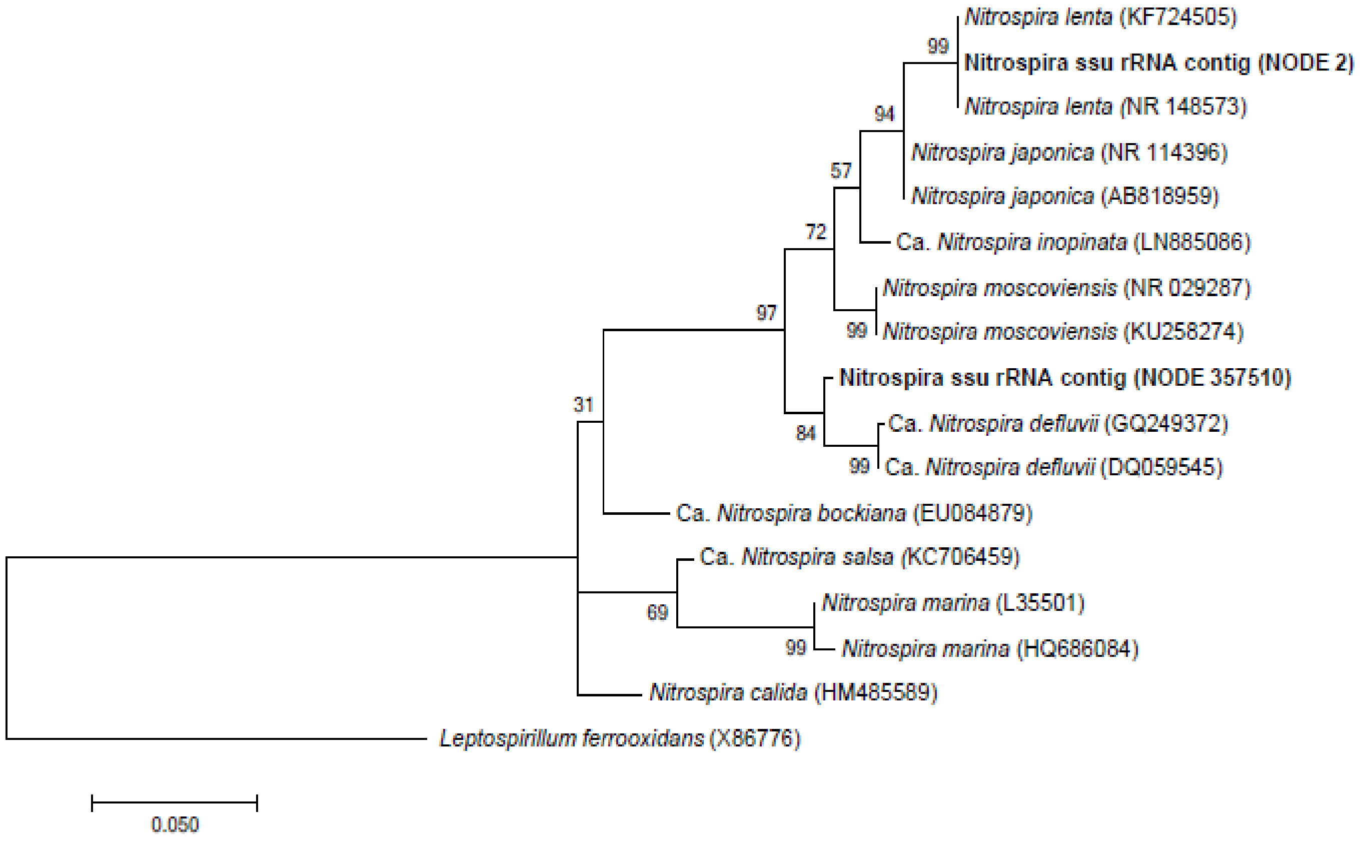


**Fig. S10**: Maximum likelihood phylogenetic tree showing the grouping of *Nitrospira* SSU rRNA contigs retrieved from metagenomic data with *Nitrospira* spp. 16S rRNA reference strains. With *Leptospirillum ferrooxidans* as the outgroup and bootstrap analysis of 1000 replicates (bootstrap values are indicated as percentages).

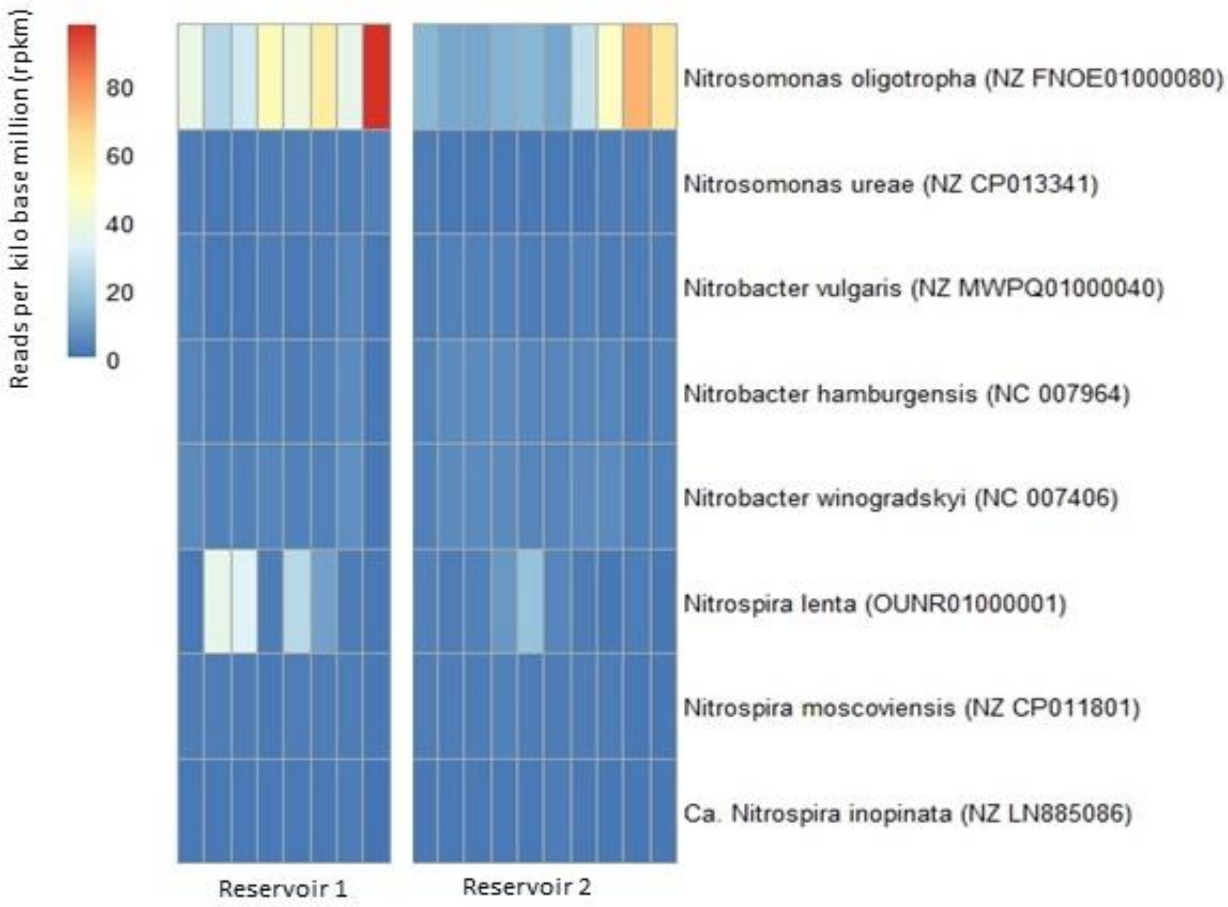


**Fig. S11**: Heatmap showing the coverage of reference genomes across all samples, which mapped on average > 1 million reads per kilo base of the reference genome (rpkm).

**Table S1A**: Chemical monitoring data for the duration of the study obtained from the utility for reservoir 1 (RES1)

| | Month | Free and available chlorine (mg/l) | Total chorine (mg/l) | Monochloramine (mg/l) | Total residual Chorine (mg/l) | Temp (°C) | Δ ammonium (mg/l) | Δ nitrite (mg/l) | Δ nitrate (mg/l) |
|---|---|---|---|---|---|---|---|---|---|
| 1 | 7-Oct-2014 | 0.53 | 2.09 | 1.56 | 2.12 | 16.1 | -0.17 | 0.00 | -0.02 |
| 2 | 4-Nov-2014 | 0.3 | 2.2 | 1.9 | 2.2 | 18.5 | 0.07 | 0.00 | 0.04 |
| 3 | 2-Dec-2014 | 0.03 | 0.74 | 0.71 | 0.81 | 22.3 | -0.02 | 0.02 | 0.06 |
| 4 | 13-Jan-2015 | 0.07 | 1.94 | 1.87 | 1.94 | 23.4 | -0.05 | 0.33 | 0.23 |
| 5 | 3-Feb-2015 | 0.11 | 2.07 | 1.96 | 2.09 | 22.7 | -0.12 | 0.29 | 0.12 |
| 6 | 3-Mar-2015 | 0.34 | 0.36 | 0.02 | 0.36 | 22.2 | -0.30 | 0.24 | 0.16 |
| 7 | 14-Apr-2015 | 0 | 0.15 | 0.15 | 0.18 | 20.8 | -0.22 | 0.36 | 0.29 |
| 8 | 5-May-2015 | 0.06 | 1.26 | 1.2 | 1.28 | 18.4 | -0.06 | 0.04 | -0.01 |
| 9 | 2-Jun-2015 | 0.06 | 0.67 | 0.61 | 0.68 | 16.8 | 0.05 | 0.13 | 0.09 |
| 10 | 7-Jul-2015 | 0.18 | 1.35 | 1.17 | 1.68 | 10.5 | -0.34 | 0.00 | 0.05 |
| 11 | 4-Aug-2015 | 0.09 | 0.96 | 0.87 | 0.95 | 12 | 0.03 | 0.00 | 0.00 |
| 12 | 1-Sep-2015 | 0.11 | 1.66 | 1.55 | 1.61 | 14.7 | 0.08 | 0.00 | 0.03 |
| 13 | 6-Oct-2015 | 0.01 | 1.42 | 1.41 | 1.42 | 19 | 0.12 | 0.00 | 0.02 |
| 14 | 3-Nov-2015 | 0.12 | 1.69 | 1.57 | 1.74 | 19.5 | 0.23 | 0.00 | -0.04 |
| 15 | 1-Dec-2015 | 0.06 | 0.11 | 0.05 | 0.13 | 20.7 | -0.04 | 0.00 | 0.01 |
| 16 | 5-Jan-2016 | 0.03 | 0.1 | 0.07 | 0.14 | 22.7 | -0.10 | 0.00 | 0.06 |
| 17 | 2-Feb-2016 | 0 | 0.33 | 0.33 | 0.35 | 23.2 | -0.08 | 0.00 | 0.00 |
| 18 | 8-Mar-2016 | 0.06 | 0.27 | 0.21 | 0.3 | 23.5 | -0.06 | 0.08 | 0.11 |
| 19 | 5-Apr-2016 | 0.03 | 0.48 | 0.45 | 0.59 | 21.7 | -0.07 | 0.00 | 0.03 |
| 20 | 10-May-2016 | 0.13 | 1.07 | 0.94 | 1.07 | 18.1 | 0.06 | 0.00 | 0.07 |
| 21 | 7-Jun-2016 | 0.24 | 0.84 | 0.6 | 0.97 | 15.2 | 0.09 | 0.00 | -0.02 |
| 22 | 5-Jul-2016 | 0.15 | 2.11 | 1.96 | 2.11 | 12.8 | - | 0.00 | 0.20 |
| 23 | 2-Aug-2016 | 0.09 | 1.84 | 1.75 | 1.99 | 11.9 | - | 0.00 | 0.02 |
| 24 | 30-Aug-2016 | 0.21 | 1.94 | 1.73 | 1.86 | 14.2 | - | 0.00 | 0.13 |

**Table S1B**: Chemical monitoring data for the duration of the study obtained from the utility for reservoir 2 (RES2)

| | Month | Free and available chlorine (mg/l) | Total chorine (mg/l) | Monochloramine (mg/l) | Total residual Chorine (mg/l) | Temp (°C) | Δ ammonium (mg/l) | Δ nitrite (mg/l) | Δ nitrate (mg/l) |
|---|---|---|---|---|---|---|---|---|---|
| 1 | 10-Oct-2014 | 0.13 | 0.64 | 0.51 | 0.98 | 17.9 | -0.15 | 0.02 | 0.08 |
| 2 | 7-Nov-2014 | 0.85 | 1.11 | 0.26 | 1.13 | 20.6 | -0.15 | -0.02 | -0.02 |
| 3 | 1-Dec-2014 | 0.04 | 1 | 0.96 | 1 | 19.3 | -0.14 | 0.05 | 0.07 |
| 4 | 16-Jan-2015 | 0.1 | 0.17 | 0.07 | 0.18 | 23 | -0.02 | -0.30 | 0.21 |
| 5 | 6-Feb-2015 | 0.04 | 0.06 | 0.02 | 0.09 | 22.8 | -0.01 | -0.60 | 0.37 |
| 6 | 6-Mar-2015 | 0.02 | 0.06 | 0.04 | 0.08 | 22.4 | -0.03 | -0.46 | 0.40 |
| 7 | 17-Apr-2015 | 0.02 | 0.05 | 0.03 | 0.07 | 21.2 | -0.02 | -0.54 | 0.39 |
| 8 | 8-May-2015 | 0.03 | 0.04 | 0.01 | 0.05 | 20 | -0.09 | -0.25 | 0.55 |
| 9 | 5-Jun-2015 | 0 | 0.02 | 0.02 | 0.03 | 16.4 | -0.06 | -0.42 | 0.07 |
| 10 | 10-Jul-2015 | 0.04 | 0.62 | 0.58 | 0.65 | 13.5 | -0.27 | 0.09 | 0.16 |
| 11 | 7-Aug-2015 | 0.2 | 1.25 | 1.05 | 1.23 | 12.9 | -0.10 | 0.00 | -0.01 |
| 12 | 4-Sep-2015 | 0.05 | 1.44 | 1.39 | 1.47 | 14.2 | -0.16 | 0.00 | 0.02 |
| 13 | 9-Oct-2015 | 0.11 | 0.82 | 0.71 | 1.21 | 18.7 | -0.21 | 0.02 | 0.08 |
| 14 | 6-Nov-2015 | 0.11 | 1.28 | 1.17 | 1.35 | 20.1 | -0.12 | 0.15 | 0.07 |
| 15 | 4-Dec-2015 | 0.12 | 1.07 | 0.95 | 1.13 | 20.5 | -0.18 | 0.28 | 0.12 |
| 16 | 8-Jan-2016 | 0.12 | 0.42 | 0.3 | 0.42 | 24 | -0.18 | 0.10 | 0.29 |
| 17 | 5-Feb-2016 | 0.11 | 0.19 | 0.08 | 0.23 | 23.4 | -0.10 | 0.06 | -0.02 |
| 18 | 11-Mar-2016 | 0.15 | 0.48 | 0.33 | 0.51 | 23.4 | -0.21 | 0.00 | 0.26 |
| 19 | 8-Apr-2016 | 0 | 0.5 | 0.5 | 0.51 | 21.6 | -0.27 | 0.00 | 0.27 |
| 20 | 13-May-2016 | 0.16 | 0.48 | 0.32 | 0.49 | 18 | -0.35 | -0.02 | 0.10 |
| 21 | 10-Jun-2016 | 0.07 | 1.55 | 1.48 | 1.57 | 15.9 | -0.34 | 0.00 | 0.22 |
| 22 | 8-Jul-2016 | 0.05 | 1.78 | 1.73 | 1.75 | 13.7 | -0.23 | 0.00 | -0.18 |
| 23 | 5-Aug-2016 | 0.12 | 1.75 | 1.63 | 1.86 | 12.3 | 0.07 | 0.00 | 0.02 |
| 24 | 2-Sep-2016 | 0.1 | 1.74 | 1.64 | 1.69 | 14.9 | -0.10 | 0.00 | -0.03 |

**Table S2**: Outline and description of the chloraminated reservoir samples collected over the 2-year study period together with the resulting DNA concentrations

| | Sample Name | Month | Year | Qubit DNA concentration (ng/ul) |
|---|---|---|---|---|
| 1 | RES1_4 | January | 2015 | 35.0 |
| 2 | RES1_5 | February | 2015 | 23.2 |
| 3 | RES1_6 | March | 2015 | 13.3 |
| 4 | RES1_17 | February | 2016 | 40.2 |
| 5 | RES1_18 | March | 2016 | 19.7 |
| 6 | RES1_19 | April | 2016 | 12.6 |
| 7 | RES1_20 | May | 2016 | 10.6 |
| 8 | RES1_21 | June | 2016 | 6.92 |
| 9 | RES2_1 | October | 2014 | 13.8 |
| 10 | RES2_5 | February | 2015 | 14.6 |
| 11 | RES2_6 | March | 2015 | 42.8 |
| 12 | RES2_7 | April | 2015 | 7.84 |
| 13 | RES2_8 | May | 2015 | 23.3 |
| 14 | RES2_10 | July | 2015 | 17.6 |
| 15 | RES2_17 | February | 2016 | 11.3 |
| 16 | RES2_18 | March | 2016 | 20.8 |
| 17 | RES2_19 | April | 2016 | 30.8 |
| 18 | RES2_20 | May | 2016 | 24.6 |

**Table S3**: The final number of reads mapped to assembly

| | Sample | Number of reads | Mapped reads to scaffolds > 500 bp | % Mapped reads |
|---|---|---|---|---|
| 1 | RES1_4 | 14 249 511 | 14 195 821 | 99.62 |
| 2 | RES1_5 | 13 019 437 | 12 929 842 | 99.31 |
| 3 | RES1_6 | 12 747 382 | 12 673 029 | 99.42 |
| 4 | RES1_17 | 14 647 502 | 14 573 146 | 99.49 |
| 5 | RES1_18 | 18 488 103 | 18 376 301 | 99.40 |
| 6 | RES1_19 | 11 184 973 | 11 116 758 | 99.39 |
| 7 | RES1_20 | 6 855 217 | 6 753 636 | 98.52 |
| 8 | RES1_21 | 16 944 215 | 16 809 348 | 99.20 |
| 9 | RES2_1 | 12 231 070 | 11 382 585 | 93.06 |
| 10 | RES2_5 | 11 677 750 | 11 487 402 | 98.37 |
| 11 | RES2_6 | 6 919 446 | 6 758 814 | 97.68 |
| 12 | RES2_7 | 11 590 760 | 11 452 070 | 98.80 |
| 13 | RES2_8 | 13 020 429 | 12 809 835 | 98.38 |
| 14 | RES2_10 | 13 802 699 | 13 303 643 | 96.38 |
| 15 | RES2_17 | 13 688 540 | 13 501 850 | 98.64 |
| 16 | RES2_18 | 13 259 038 | 13 173 737 | 99.36 |
| 17 | RES2_19 | 21 940 278 | 21 877 986 | 99.72 |
| 18 | RES2_20 | 12 542 819 | 12 483 217 | 99.52 |

**Table S4**: Summary of the genes identified (including KEGG orthology numbers) involved in nitrogen transformation pathways in chloraminated reservoir samples and their cumulative coverage (rpkm) in each reservoir

| **K number** | **Enzyme** | **Enzyme name** | **Reaction** | **Process** | **RES1*** | **RES2*** |
|---|---|---|---|---|---|---|
| K10944 | *amoA* | Ammonia monooxygenase, subunit A | $NH_4^+ + O_2 + 2e^- \text{-->} NH_2OH + H_2O$ | Ammonia oxidation - Nitrification | 1143.07 | 1671.50 |
| K10945 | *amoB* | Ammonia monooxygenase, subunit B | $NH_4^+ + O_2 + 2e^- \text{-->} NH_2OH + H_2O$ | Ammonia oxidation - Nitrification | 2922.21 | 2165.83 |
| K10946 | *amoC* | Ammonia monooxygenase, subunit C | $NH_4^+ + O_2 + 2e^- \text{-->} NH_2OH + H_2O$ | Ammonia oxidation - Nitrification | 4625.20 | 4807.07 |
| K10535 | *hao* | Hydroxylamine oxidoreductase | $NH_2OH \text{-->} NO + 3e^- + 3H^+$ | Hydroxylamine oxidation - Nitrification | 2486.24 | 2169.57 |
| K05601 | *hcp* | Hydroxylamine reductase | $NH_2OH + H_2O \text{-->} NH_4^+ + O_2 + 2e^-$ | Hydroxylamine reduction - Nitrification | 6.76 | 53.03 |
| K00370 | *nxrA* | Nitrite oxidoreductase, alpha subunit | $NO_2^- + H_2O \text{-->} NO_3^- + 2e^- + 2H^+$ | Nitrite oxidation - Nitrification | 346.72 | 183.14 |
| K00371 | *nxrB* | Nitrite oxidoreductase, beta subunit | $NO_2^- + H_2O \text{-->} NO_3^- + 2e^- + 2H^+$ | Nitrite oxidation - Nitrification | 335.18 | 135.21 |
| K00370 | *narG* | Respiratory nitrate reductase, alpha subunit | $NO_3^- + 2e^- + 2H^+ \text{-->} NO_2^- + H_2O$ | Dissimilatory nitrate reduction | 66.10 | 278.95 |
| K00371 | *narH* | Respiratory nitrate reductase, beta subunit | $NO_3^- + 2e^- + 2H^+ \text{-->} NO_2^- + H_2O$ | Dissimilatory nitrate reduction | 35.15 | 205.32 |
| K00373 | *narJ* | Respiratory nitrate reductase, delta subunit (molybdenum cofactor assembly chaperone) | $NO_3^- + 2e^- + 2H^+ \text{-->} NO_2^- + H_2O$ | Dissimilatory nitrate reduction | 18.09 | 159.90 |
| K00374 | *narI* | Respiratory nitrate reductase, gamma subunit | $NO_3^- + 2e^- + 2H^+ \text{-->} NO_2^- + H_2O$ | Dissimilatory nitrate reduction | 23.94 | 182.24 |
| K00372 | *nasA* | Assimilatory nitrate reductase, large catalytic subunit | $NO_3^- + 2e^- + 2H^+ \text{-->} NO_2^- + H_2O$ | Assimilatory nitrate reduction | 455.05 | 701.03 |
| K00362 | *nirB* | Nitrite reductase (NAD(P)H), large subunit | $NO_2^- + 6e^- + 8H^+ \text{-->} NH_4^+ + 2H_2O$ | Dissimilatory nitrite reduction | 248.74 | 437.86 |
| K00363 | *nirD* | Nitrite reductase (NAD(P)H), small subunit | $NO_2^- + 6e^- + 8H^+ \text{-->} NH_4^+ + 2H_2O$ | Dissimilatory nitrite reduction | 94.70 | 310.78 |
| K03385 | *nrfA* | Cytochrome c-552 | $NO_2^- + 6e^- + 8H^+ \text{-->} NH_4^+ + 2H_2O$ | Dissimilatory nitrite reduction | 3.98 | 32.21 |
| K00366 | *nirA* | Ferredoxin-nitrite reductase | $NO_2^- + 6e^- + 8H^+ \text{-->} NH_4^+ + 2H_2O$ | Assimilatory nitrite reduction | 729.37 | 692.10 |
| K00368 | *nirK* | Nitrite reductase (NO-forming) | $NO_2^- + e^- + 2H^+ \text{-->} NO + H_2O$ | Nitrite reduction - Nitric oxide (NO) forming | 2016.40 | 2043.11 |
| K02448 | *norD* | Nitric oxide reductase, D protein | $2NO + 2e^- + 2H^+ \text{-->} N_2O + H_2O$ | Nitric oxide reduction | 856.37 | 699.77 |
| K04561 | *norB* | Nitric oxide reductase, subunit B | $2NO + 2e^- + 2H^+ \text{-->} N_2O + H_2O$ | Nitric oxide reduction | 1363.59 | 971.78 |

| | | | | | | |
|---|---|---|---|---|---|---|
| K02305 | *norC* | Nitric oxide reductase, subunit C | $2NO + 2e^- + 2H^+ \dashrightarrow N_2O + H_2O$ | Nitric oxide reduction | 1060.40 | 427.07 |
| K04748 | *norQ* | Nitric oxide reductase, Q protein | $2NO + 2e^- + 2H^+ \dashrightarrow N_2O + H_2O$ | Nitrous-oxide reduction | 1121.85 | 996.92 |
| K00376 | *nosZ* | Nitrous-oxide reductase | $N_2O + 2e^- 2H^+ \dashrightarrow N_2 + H_2O$ | Nitrogenase - Nitrogen fixation | 62.22 | 85.89 |
| K02586 | *nifD* | Nitrogenase molybdenum-iron protein, alpha chain | $N_2 + 8e^- + 8H^+ + 16ATP \dashrightarrow 2NH_3 + H_2 + 16ADP + 16P_i$ | Nitrogenase - Nitrogen fixation | 32.06 | 8.58 |
| K02591 | *nifK* | Nitrogenase molybdenum-iron protein, beta chain | $N_2 + 8e^- + 8H^+ + 16ATP \dashrightarrow 2NH_3 + H_2 + 16ADP + 16P_i$ | Nitrogenase - Nitrogen fixation | 30.73 | 4.64 |
| K02588 | *nifH* | Nitrogenase iron protein | $N_2 + 8e^- + 8H^+ + 16ATP \dashrightarrow 2NH_3 + H_2 + 16ADP + 16P_i$ | Nitrogenase - Nitrogen fixation | 31.56 | 9.82 |

* Cumulative reads per kilobase million (rpkm) of each gene in each reservoir

**Table S5**: Reference genomes of nitrifying organisms

| | Reference name | Accession number | Nitrifier type |
|---|---|---|---|
| 1 | *Ca*. Nitrosoarchaeum sp. | NSIR01000002 | Ammonia oxidizing archaea |
| 2 | *Ca*. Nitrosoarchaeum koreensis | NZ AFPU01000001 | Ammonia oxidizing archaea |
| 3 | *Ca*. Nitrosopumilus adriaticus | NZ CP011070 | Ammonia oxidizing archaea |
| 4 | *Ca*. Nitrosopumilus koreensis | CP003842 | Ammonia oxidizing archaea |
| 5 | *Ca*. Nitrosopumilus piranensis | CP010868 | Ammonia oxidizing archaea |
| 6 | *Ca*. Nitrosopumilus sp. | NC 018656 | Ammonia oxidizing archaea |
| 7 | *Ca*. Nitrososphaera evergladensis | CP007174 | Ammonia oxidizing archaea |
| 8 | *Ca*. Nitrososphaera gargensis | CP002408 | Ammonia oxidizing archaea |
| 9 | *Cenarchaeum symbiosum* | DP000238 | Ammonia oxidizing archaea |
| 10 | *Nitrosopumilus maritimus* | CP000866 | Ammonia oxidizing archaea |
| 11 | *Nitrososphaera viennensis* | NZ CP007536 | Ammonia oxidizing archaea |
| 12 | *Nitrosococcus halophilus* | NC 013960 | Ammonia oxidizing bacteria |
| 13 | *Nitrosococcus oceani* | NC 007484 | Ammonia oxidizing bacteria |
| 14 | *Nitrosococcus watsonii* | NC 014315 | Ammonia oxidizing bacteria |
| 15 | *Nitrosomonas aestuarii* | NZ FOSP01000116 | Ammonia oxidizing bacteria |
| 16 | *Nitrosomonas communis* | NZ CP011451 | Ammonia oxidizing bacteria |
| 17 | *Nitrosomonas cryotolerans* | NZ FSRO01000002 | Ammonia oxidizing bacteria |
| 18 | *Nitrosomonas europaea* | NC 004757 | Ammonia oxidizing bacteria |
| 19 | *Nitrosomonas eutropha* | NC 008344 | Ammonia oxidizing bacteria |
| 20 | *Nitrosomonas halophila* | NZ FNOY01000156 | Ammonia oxidizing bacteria |
| 21 | *Nitrosomonas marina* | NZ FOCP01000068 | Ammonia oxidizing bacteria |
| 22 | *Nitrosomonas mobilis* | NZ FMWO01000108 | Ammonia oxidizing bacteria |
| 23 | *Nitrosomonas nitrosa* | NZ FOUF01000076 | Ammonia oxidizing bacteria |
| 24 | *Nitrosomonas oligotropha* | NZ FNOE01000080 | Ammonia oxidizing bacteria |
| 25 | *Nitrosomonas ureae* | NZ CP013341 | Ammonia oxidizing bacteria |
| 26 | *Nitrosospira briensis* | NZ CP012371 | Ammonia oxidizing bacteria |
| 27 | *Nitrosospira lacus* | NZ CP021106 | Ammonia oxidizing bacteria |
| 28 | *Nitrosospira multiformis* | NC 007614 | Ammonia oxidizing bacteria |
| 29 | *Ca*. Nitrospira inopinata | NZ LN885086 | Comammox bacteria |
| 30 | *Ca*. Nitrospira nitrificans | NZ CZPZ01000001 | Comammox bacteria |
| 31 | *Ca*. Nitrospira nitrosa | NZ CZQA01000001 | Comammox bacteria |
| 32 | *Nitrospira* sp. | LNDU01000039 | Comammox bacteria |
| 33 | *Ca*. Nitrospira defluvii | NC 014355 | Nitrite oxidizing bacteria |
| 34 | *Nitrobacter vulgaris* | NZ MWPQ01000040 | Nitrite oxidizing bacteria |
| 35 | *Nitrobacter hamburgensis* | NC 007964 | Ammonia oxidizing bacteria |
| 36 | *Nitrobacter winogradskyi* | NC 007406 | Ammonia oxidizing bacteria |
| 37 | *Nitrospira japonica* | NZ LT828648 | Nitrite oxidizing bacteria |
| 38 | *Nitrospira lenta* | OUNR01000001 | Nitrite oxidizing bacteria |

| | | | |
|---|---|---|---|
| 39 | *Nitrospira moscoviensis* | NZ CP011801 | Nitrite oxidizing bacteria |
| 40 | *Ca*. Brocadia caroliniensis | AYTS01000035 | Anammox bacteria |
| 41 | *Ca*. Brocadia fulgida | LAQJ01000121 | Anammox bacteria |
| 42 | *Ca*. Brocadia sapporoensis | NZ MJUW02000122 | Anammox bacteria |
| 43 | *Ca*. Jettenia caeni | NZ BAFH01000003 | Anammox bacteria |
| 44 | *Ca*. Kuenenia stuttgartiensis | NZ LT934425 | Anammox bacteria |

**Table S6**: Characteristics of constructed Metagenome Assembled Genomes (MAGs)

| MAG | Taxonomy[a] | MiGA | Completeness % | Contamination % | Quality % | G+C content % | Number of contigs | Genetic size (Mb) | Largest contig (bp) | N50 | Number of ORF's | Number of genes annotated | Coverage RES1 (rpkm)[c] | Coverage RES2 (rpkm)[c] |
|---|---|---|---|---|---|---|---|---|---|---|---|---|---|---|
| c2 | *Bacteroidetes Cytophagia Cytophagale Flammeovirgaceae* | *Marivirga tractuosa* (50.15% AAI) | 91.9 | 2.7 | 78.4 | 41.79 | 396 | 3.87 | 82 589 | 14 095 | 3747 | 1452 (38.8%) | 0.1 ± 0.07 | 0.4 ± 0.46 |
| c3 | *Alphaproteobacteria Rhodospirillales Acetobacteraceae* | *Roseomonas* sp. (56.54% AAI) | 94.6 | 0.9 | 90.1 | 69.05 | 292 | 4.52 | 125 874 | 24 612 | 4612 | 2024 (43.9%) | 1.32 ± 2.46 | 1.35 ± 1.71 |
| c4 | *Alphaproteobacteria Caulobacterales Caulobacteraceae* | *Caulobacteraceae* bacterium (52.29% AAI) | 91 | 0 | 91 | 67.51 | 163 | 2.72 | 83 313 | 25 390 | 2817 | 1389 (49.3%) | 0.79 ± 1.16 | 1.63 ± 1.3 |
| c14 | *Gammaprotebacteria* | *Thiohalobacter thiocyanaticus* (48.24% AAI) | 90.1 | 1.8 | 81.1 | 67.88 | 729 | 5.51 | 34 030 | 9 353 | 5556 | 2325 (41.8%) | 0.14 ± 0.1 | 0.56 ± 0.53 |
| c19 | *Alphaproteobacteria Sphingomonadales Erythrobacteraceae Porphyrobacter* | *Porphyrobacter* sp. CACIAM (81.5 ± 14,4% AAI) | 94.6 | 0.9 | 90.1 | 64.06 | 58 | 3.41 | 476 516 | 264 474 | 3279 | 1586 (48.4%) | 0.05 ± 0.01 | 4.6 ± 7.23 |
| c24 | *Betaproteobacteria Burkholderiales* | *Rhizobacter gummiphilus* (62.69% AAI) | 95.5 | 1.8 | 86.5 | 64.87 | 64 | 3.46 | 232 861 | 99 668 | 3260 | 1832 (56.2%) | 0.09 ± 0.06 | 0.49 ± 0.53 |
| c26 | *Betaproteobacteria Methylophilales Methylophilaceae Methylotenera* | *Methylotenera mobilis* (62.03% AAI) | 95.5 | 1.8 | 86.5 | 41.94 | 12 | 2.03 | 735 558 | 469 317 | 2016 | 1268 (62.9%) | 0.02 ± 0.01 | 0.58 ± 1.17 |
| c27 | *Alphaproteobacteria Rhizobiales Bradyrhizobiaceae Bosea* | *Bosea* sp. PAMC (79.03 ± 15.87% AAI) | 95.5 | 0.9 | 91 | 66.75 | 117 | 4.81 | 204 440 | 72 057 | 4670 | 2376 (50.9%) | 0.06 ± 0.02 | 2.6 ± 3.8 |
| c31 | *Alphaproteobacteria Rhizobiales Beijerinckiaceae* | *Methylocella silvestris* (56.71% AAI) | 95.5 | 1.8 | 86.5 | 62.55 | 58 | 4.43 | 399 037 | 106 052 | 4266 | 1942 (45.5%) | 0.08 ± 0.06 | 0.78 ± 1.06 |
| c33 | *Alphaproteobacteria Rhizobiales Bradyrhizolbiaceae Oligotropha* | *Oligotropha carboxidovorans* (83.31 ± 12.94% AAI) | 94.6 | 1.8 | 85.6 | 61.68 | 51 | 3.45 | 253 596 | 122 000 | 3301 | 1794 (54.3%) | 0.05 ± 0.03 | 0.55 ± 0.82 |
| c34 | *Gemmatimonadetes Gemmatimonadales Gemmatimonadaceae Gemmatirosa* | *Gemmatimonas phototrophica* (53.28% AAI) | 73 | 1.8 | 64 | 67.83 | 537 | 2.74 | 26 761 | 5 888 | 2892 | 1226 (42.4%) | 0.16 ± 0.08 | 0.51 ± 0.47 |
| c35 | *Alphaproteobacteria Rhizobiales* | *Hyphomicrobium denitrificans* | 88.3 | 1.8 | 79.3 | 58.89 | 431 | 4.00 | 85 476 | 13 110 | 4239 | 1835 (43.3%) | 2.3 ± 2.22 | 1.12 ± 1 |

| | | | | | | | | | | | | | |
|---|---|---|---|---|---|---|---|---|---|---|---|---|---|
| | *Hyphomicrobiaceae*<br>*Hyphomicrobium* | (87.12 ±14.44% AAI) | | | | | | | | | | | |
| c38 | *Alphaproteobacteria*<br>*Sphingomonadales*<br>*Sphingomonadaceae*<br>*Novosphingobium* | *Novosphingobium*<br>*aromaticivorans*<br>(66.34% AAI) | 93.7 | 0.9 | 89.2 | 68.83 | 64 | 4.07 | 251 248 | 119 692 | 3727 | 1770<br>(47.5%) | 0.07 ± 0.07 | 0.53 ±<br>0.54 |
| c39 | *Betaproteobacteria*<br>*Sulfuricellales*<br>*Sulfuricellaceae*<br>*Sulfuricella* | *Sulfuricella*<br>*denitrificans*<br>(84.31 ± 17.04% AAI) | 94.6 | 3.6 | 76.6 | 58.61 | 92 | 3.57 | 180 696 | 66 989 | 3485 | 1851<br>(53.1%) | 0.07 ± 0.02 | 0.5 ± 1.02 |
| c41 | *Alphaproteobacteria*<br>*Sphingomonadales*<br>*Erythrobacteraceae*<br>*Porphyrobacter* | *Porphyrobacter* sp.<br>LM 6<br>(97.28 ± 7.93% ANI) | 93.7 | 1.8 | 84.7 | 64.72 | 33 | 3.04 | 397 646 | 156 172 | 2951 | 1471<br>(49.8%) | 0.17 ± 0.1 | 3.74 ± 8.4 |
| c43 | *Alphaproteobacteria*<br>*Rhodospirillales* | *Azospirillum*<br>*brasilense*<br>(49.02% AAI) | 93.7 | 2.7 | 80.2 | 66.19 | 611 | 5.89 | 51 990 | 13 561 | 6212 | 2683<br>(43.2%) | 0.5 ± 0.56 | 0.38 ±<br>0.38 |
| c47 | *Betaproteobacteria*<br>*Nitrosomonadales*<br>*Gallionellaceae*<br>*Sideroxydans* | *Sideroxydans*<br>*lithotrophicus*<br>(77.27 ± 16,96% AAI) | 73 | 2.7 | 59.5 | 55.31 | 455 | 2.29 | 26 046 | 5 807 | 2655 | 1512<br>(56.9%) | 0.3 ± 0.35 | 0.21 ±<br>0.21 |
| c48.1[a] | *Alphaproteobacteria*<br>*Sphingomonadales*<br>*Sphingomonadaceae*<br>*Sphingopyxis* | *Sphingopyxis*<br>*terrae*<br>(63.28% AAI) | 94.6 | 1.8 | 85.6 | 63.12 | 29 | 2.95 | 433 016 | 176 469 | 2853 | 1429<br>(50.1%) | 0.06 ± 0.03 | 1.85 ±<br>3.71 |
| c48.2[a] | *Alphaproteobacteria*<br>*Rhizobiales*<br>*Bradyrhizobiaceae* | *Bradyrhizobium*<br>*lablabi*<br>(64.55% AAI) | 94.6 | 0.9 | 90.1 | 62.2 | 16 | 4.57 | 902 226 | 479 955 | 4332 | 1996<br>(46.1%) | 0.04 ± 0.01 | 1.68 ±<br>2.22 |
| c49 | *Alphaproteobacteria*<br>*Rhizobiales* | *Rhizobium*<br>*tropici*<br>(45.25% AAI) | 87.4 | 0.9 | 82.9 | 69.43 | 978 | 4.80 | 38 978 | 5 604 | 5423 | 2260<br>(41.7%) | 0.23 ± 0.28 | 0.6 ± 0.68 |
| c51 | *Nitrospira*<br>*Nitrospirales*<br>*Nitrospiraceae*<br>*Nitrospira* | *Nitrospira*<br>*moscoviensis*<br>(62.69% AAI) | 92.8 | 5.4 | 65.8 | 57.78 | 80 | 4.27 | 1 125<br>973 | 247 953 | 4354 | 1608<br>(36.9%) | 16.93 ±<br>20.05 | 4.36 ±<br>6.92 |
| c56 | *Nitrospira*<br>*Nitrospirales*<br>*Nitrospiraceae*<br>*Nitrospira* | *Nitrospira*<br>*moscoviensis*<br>(62.1% AAI) | 77.5 | 1.8 | 68.5 | 57.14 | 443 | 3.69 | 44 036 | 11 519 | 4121 | 1498<br>(36.4%) | 0.43 ± 0.5 | 0.23 ±<br>0.17 |
| c58 | *Betaproteobacteria*<br>*Nitrosomonadales*<br>*Nitrosomonadaceae*<br>*Nitrosomonas* | *Nitrosomonas* sp.<br>Is79A3<br>(66.34% AAI) | 94.6 | 0.9 | 90.1 | 48.69 | 70 | 3.12 | 237 347 | 90 243 | 2951 | 1486<br>(50.4%) | 7.12 ±<br>10.39 | 22.12 ±<br>27.3 |
| c59 | *Alphaproteobacteria*<br>*Rhodobacterales*<br>*Rhodobacteraceae* | *Rhodobacter* sp.<br>CZR27<br>(62.45% AAI) | 93.7 | 1.8 | 84.7 | 66.06 | 359 | 4.33 | 103 413 | 17 991 | 4620 | 2192<br>(47.4%) | 0.14 ± 0.08 | 0.51 ±<br>0.85 |
| c60 | *Betaproteobacteria* | *Methylotenera* | 95.5 | 1.8 | 86.5 | 45.61 | 24 | 2.53 | 486 922 | 217 332 | 2401 | 1424 | 0.02 ± 0.01 | 2.44 ± |

| | | | | | | | | | | | | | | |
|---|---|---|---|---|---|---|---|---|---|---|---|---|---|---|
| | *Nitrosomonadales*<br>*Methylophilaceae*<br>*Methylotenera* | *mobilis*<br>(77.35 ± 15,15% AAI) | | | | | | | | | | (59.3%) | | 4.84 |
| c61 | *Betaproteobacteria*<br>*Nitrosomonadales*<br>*Gallionellaceae*<br>*Sideroxydans/Gallionella* | *Sideroxydans*<br>*lithotrophicus*<br>(67.0% AAI) | 94.6 | 0.9 | 90.1 | 56.92 | 50 | 2.75 | 341 276 | 100 797 | 2814 | 1477<br>(52.5%) | 1.65 ± 1.82 | 0.09 ± 0.1 |
| c64 | *Alphaproteobacteria*<br>*Caulobacterales* | *Caulobacteraceae*<br>bacterium<br>(52.17% AAI) | 87.4 | 0 | 87.4 | 65.98 | 313 | 2.43 | 54 901 | 10 673 | 2708 | 1354<br>(50.0%) | 0.8 ± 0.87 | 0.27 ±<br>0.26 |
| c65 | *Alphaproteobacteria*<br>*Sphingomonadales*<br>*Sphingomonadaceae* | *Sphingomonas*<br>*panacis*<br>(61.83% AAI) | 93.7 | 0 | 93.7 | 61.44 | 33 | 3.23 | 571 943 | 183 848 | 3132 | 1541<br>(49.2%) | 0.99 ± 1.73 | 1.18 ± 2.5 |
| c69 | *Betaproteobacteria*<br>*Nitrosomonadales*<br>*Nitrosomonadaceae* | *Nitrosospira*<br>*lacus*<br>(52.37% AAI) | 93.7 | 1.8 | 84.7 | 60.83 | 82 | 3.91 | 228 320 | 102 579 | 3874 | 1967<br>(50.8%) | 0.05 ± 0.03 | 2.75 ±<br>4.57 |
| c70 | *Alphaproteobacteria*<br>*Sphingomonadales*<br>*Sphingomonadaceae*<br>*Sphingomonas* | *Sphingomonas*<br>*panacis*<br>(61.38% AAI) | 94.6 | 3.6 | 76.6 | 63.51 | 44 | 3.14 | 382 616 | 247 649 | 3082 | 1600<br>(51.9%) | 10.74 ±<br>18.11 | 11.95 ±<br>17.62 |
| c72 | *Alphaproteobacteria*<br>*Sphingomonadales*<br>*Sphingomonadaceae*<br>*Sphingomonas* | *Sphingomonas*<br>*panacis*<br>(62.63% AAI) | 93.7 | 0 | 93.7 | 61.81 | 31 | 3.89 | 611 207 | 391 571 | 3818 | 1682<br>(44.1%) | 2.76 ± 3.5 | 2.84 ±<br>5.13 |
| c74 | *Alphaproteobacteria*<br>*Rhizobiales*<br>*Hyphomicrobiaceae* | *Hyphomicrobium*<br>*nitrativorans*<br>(54.07% AAI) | 92.8 | 0.9 | 88.3 | 65.43 | 103 | 5.94 | 294 341 | 93 246 | 5453 | 2251<br>(41.3%) | 0.33 ± 0.72 | 1.07 ±<br>1.41 |
| c76 | *Betaproteobacteria*<br>*Nitrosomonadales*<br>*Gallionellaceae*<br>*Gallionella* | *Gallionella*<br>*capsiferriformans*<br>(79.91 ± 16.11% AAI) | 92.8 | 0.9 | 88.3 | 53.15 | 130 | 3.13 | 121 681 | 37 955 | 3102 | 1586<br>(51.1%) | 0.05 ± 0.02 | 0.34 ±<br>0.69 |
| c77 | *Betaproteobacteria*<br>*Nitrosomonadales*<br>*Methylophilaceae*<br>*Methylotenera* | *Methylotenera*<br>*versatilis*<br>(64.03% AAI) | 95.5 | 2.7 | 82 | 42.38 | 11 | 2.36 | 828 080 | 282 428 | 2257 | 1324<br>(58.7%) | 0.03 ± 0.03 | 0.62 ±<br>1.76 |
| c83 | *Betaproteobacteria*<br>*Nitrosomonadales*<br>*Gallionellaceae*<br>*Sideroxydans/Gallionella* | *Gallionella*<br>*capsiferriformans*<br>(62.29% AAI) | 95.5 | 0.9 | 91 | 56.01 | 132 | 3.01 | 195 785 | 37 811 | 2977 | 1581<br>(53.1%) | 0.05 ± 0.03 | 0.6 ± 1.5 |
| c86 | *Alphaproteobacteria*<br>*Rhizobiales* | *Rhizobium* sp.<br>N324<br>(47.54% AAI) | 81.1 | 5.4 | 54.1 | 63.81 | 669 | 3.19 | 45 151 | 5 479 | 3618 | 1658<br>(45.8%) | 0.2 ± 0.23 | 0.52 ± 0.6 |
| c90 | *Alphaproteobacteria*<br>*Rhizobiales*<br>*Hyphomicrobiaceae* | *Hyphomicrobium*<br>*nitrativorans*<br>(54.02% AAI) | 93.7 | 0.9 | 89.2 | 63.9 | 87 | 5.15 | 412 933 | 130 872 | 4854 | 2111<br>(43.5%) | 4.07 ± 2.22 | 2.77 ±<br>0.97 |
| c93 | *Alphaproteobacteria*<br>*Rhizobiales* | *Methylobacterium* sp.<br>C1 | 94.6 | 3.6 | 76.6 | 66.32 | 231 | 5.46 | 316 841 | 51 541 | 5451 | 2106<br>(38.6%) | 1.93 ± 3.36 | 2.57 ± 4.3 |

| | | | | | | | | | | | | | |
|---|---|---|---|---|---|---|---|---|---|---|---|---|---|
| | *Methylobacteriaceae*<br>*Methylobacterium* | (65.19% AAI) | | | | | | | | | | | |
| c94 | *Planctomycetes*<br>*Planctomycetia*<br>*Planctomycetales*<br>*Planctomycetaceae* | *Singulisphaera acidiphila*<br>(41.76% AAI) | 91.9 | 2.7 | 78.4 | 46.18 | 124 | 4.80 | 525 508 | 101 160 | 3712 | 1329<br>(35.9%) | 0.12 ± 0.08 | 0.44 ± 1.02 |
| c97 | Unknown<br>*Planctomycetes*<br>*Phycisphaerae* | *Phycisphaera mikurensis*<br>(40.47% AAI) | 89.2 | 0 | 89.2 | 65.03 | 76 | 3.60 | 258 711 | 82 719 | 2976 | 1133<br>(38.1%) | 0.03 ± 0.02 | 0.68 ± 0.92 |
| c102 | *Betaproteobacteria*<br>*Nitrosomonadales*<br>*Gallionellaceae*<br>*Sideroxydans/Gallionella* | *Sideroxydans lithotrophicus*<br>(66.35% AAI) | 93.7 | 1.8 | 84.7 | 55.38 | 116 | 2.63 | 140 207 | 39 153 | 2675 | 1520<br>(56.8%) | 0.16 ± 0.07 | 0.23 ± 0.42 |
| c103.1[a] | *Alphaproteobacteria*<br>*Rhizobiales*<br>*Hyphomicrobiaceae*<br>*Hyphomicrobium* | *Hyphomicrobium denitrificans*<br>(61.59% AAI) | 95.5 | 7.2 | 59.5 | 59.99 | 12 | 3.65 | 1 305 095 | 829 204 | 3405 | 1705<br>(50.1%) | 1.92 ± 1.4 | 2.01 ± 1.75 |
| c103.2[a] | *Alphaproteobacteria*<br>*Rhizobiales* | *Chelatococcus* sp.<br>(51.55% AAI) | 94.6 | 0.9 | 90.1 | 60.73 | 67 | 3.28 | 248 172 | 102 576 | 3179 | 1765<br>(55.5%) | 24.54 ± 12.15 | 19.91 ± 10.14 |
| c104 | *Alphaproteobacteria*<br>*Sphingomonadales*<br>*Sphingomonadaceae* | *Sphingomonas panacis*<br>(61.76% AAI) | 94.6 | 0.9 | 90.1 | 66.91 | 96 | 3.25 | 161 741 | 54 966 | 3263 | 1554<br>(47.6%) | 4.91 ± 6.11 | 3.59 ± 2.98 |
| c107 | *Betaproteobacteria*<br>*Nitrosomonadales*<br>*Nitrosomonadaceae*<br>*Nitrosomonas* | *Nitrosomonas* sp. Is79A3<br>(66.17% AAI) | 93.7 | 4.5 | 71.2 | 48.3 | 163 | 3.66 | 208 136 | 64 726 | 3461 | 1687<br>(48.7%) | 42.54 ± 26.31 | 11.82 ± 6.23 |
| c109 | *Betaproteobacteria*<br>*Nitrosomonadales*<br>*Nitrosomonadaceae* | *Nitrosospira* lacus<br>(52.47% AAI) | 95.5 | 0.9 | 91 | 62.68 | 59 | 3.80 | 299 525 | 118 187 | 3636 | 1832<br>(50.4%) | 0.33 ± 0.27 | 1.78 ± 4.76 |
| c114 | *Betaproteobacteria*<br>*Burkholderiales*<br>*Comamonadaceae* | *Acidovorax* sp. NA3<br>(65.46% AAI) | 91.9 | 2.7 | 78.4 | 63.89 | 497 | 3.99 | 114 432 | 12 659 | 4261 | 2171<br>(51.0%) | 0.14 ± 0.06 | 0.91 ± 2.04 |

[a] Manually curated Metagenome Assembled Genomes (MAGs)

[b] Taxon was assigned when at least 75% of the identified genes resulted in a concordant taxonomy.

[c] Values calculated as the mean coverage of each MAG across all sample within each reservoir (Mean ± Standard deviation)

**Table S7**: Percentage coverage of SSU rRNA contigs identified as bacterial phyla, eukaryota and unclassified averaged across both reservoirs (coverage calculated for contigs > 250bp)

| | | | RES1 | | RES2 | |
|---|---|---|---|---|---|---|
| | | | MRA | SD | MRA | SD |
| Bacterial phyla | *Proteobacteria* | | 81.60 | 13.32 | 82.98 | 10.28 |
| | | *Alphaproteobacteria* | 42.99 | 13.97 | 52.75 | 15.36 |
| | | *Deltaproteobacteria* | 0.08 | 0.08 | 0.09 | 0.09 |
| | | Total *Gammaproteobacteria* | 38.53 | 14.54 | 30.15 | 17.02 |
| | | *Gammaproteobacteria, Betaproteobacteriales* | 38.05 | 14.47 | 29.46 | 17.14 |
| | | Other *Gammaproteobacteria* | 0.48 | 0.37 | 0.69 | 0.35 |
| | *Nitrospirae* | | 11.25 | 13.48 | 2.28 | 3.83 |
| | *Bacteroidetes* | | 0.40 | 0.33 | 0.80 | 0.88 |
| | *Planctomycetes* | | 0.16 | 0.14 | 0.58 | 0.94 |
| | *Actinobacteria* | | 0.43 | 0.26 | 0.30 | 0.29 |
| | *Acidobacteria* | | 0.12 | 0.12 | 0.27 | 0.28 |
| | *Gemmatimonadetes* | | 0.19 | 0.18 | 0.10 | 0.16 |
| | *Patescibacteria* | | 0.11 | 0.10 | 0.08 | 0.10 |
| | *Verrucomicrobia* | | 0.04 | 0.07 | 0.10 | 0.10 |
| | *Cyanobacteria* | | 0.02 | 0.03 | 0.09 | 0.11 |
| | *Spirochaetes* | | 0.03 | 0.06 | 0.06 | 0.11 |
| | *Chloroflexi* | | 0.03 | 0.07 | 0.05 | 0.06 |
| | *Chlamydiae* | | 0.03 | 0.05 | 0.02 | 0.03 |
| Eukaryota | | | 2.05 | 1.26 | 4.64 | 5.55 |
| Unclassified contigs | | | 3.55 | 1.17 | 7.58 | 3.66 |

* MRA – Mean relative abunndance
SD – Standard deviation

**Table S8**: Percentage coverage of SSU rRNA contigs identified as Eukaryota phyla averaged across both reservoirs (coverage calculated for contigs > 250bp)

| | RES1 | | RES2 | |
|---|---|---|---|---|
| | MRA | SD | MRA | SD |
| *Metazoa* | 0.20 | 0.14 | 1.12 | 1.79 |
| Unclassified Eukaryota | 0.31 | 0.08 | 0.92 | 1.24 |
| *Opisthokonta* | 0.61 | 1.29 | 0.49 | 1.01 |
| *Ochrophyta* | 0.67 | 0.65 | 0.39 | 0.69 |
| *Stramenopiles* | 0.01 | 0.00 | 0.16 | 0.33 |
| *Amoebozoa* | 0.03 | 0.09 | 0.06 | 0.11 |
| *Bacillariophyta* | 0.01 | 0.02 | 0.05 | 0.04 |
| *Viridiplantae* | 0.03 | 0.08 | 0.04 | 0.07 |

* MRA – Mean relative abunndance
SD – Standard deviation
